# Neurological Disorder after Severe Pneumonia is Associated with Translocation of Bacteria from Lung to Brain

**DOI:** 10.1101/2022.12.30.522351

**Authors:** Qingle Ma, Chenlu Yao, Yi Wu, Heng Wang, Qin Fan, Qianyu Yang, Jialu Xu, Huaxing Dai, Yue Zhang, Fang Xu, Ting Lu, Chao Wang

**Author notes:** Corresponding author: Chao Wang.

## Abstract

The neurological disorder is a common feature in patients who recovered from severe acute pneumonia. However, the underline mechanisms remain not very clear. Here we show that these neurological syndromes after severe acute pneumonia are partly attributed to the translocation of bacteria from the lung to the brain during pneumonia. We detected an emerging and increased bacteria in the brain tissue of mice with lipopolysaccharide-induced experimental severe pneumonia. Interestingly, using 16S rDNA amplification sequencing, similarities were found between the brain’s flora species and those of the lungs, indicating the bacteria in the brain may originate from the lung. We also observed the impairment of the lung-blood barrier and brain-blood barrier, simultaneously, allowing lung bacteria invade the brain during pneumonia. An elevated microglia and astrocytes activation signature through bacterial infection-related pathways is observed by single-cell RNA sequencing, indicating a bacteria-induced disruption of brain homeostasis. Rapamycin delivered by platelet-derived extracellular vesicles provides an effective strategy to rescue the dysfunction of microglia and astrocytes, and relief neurological disorders. Collectively, we identify lung bacteria that play a role in altering brain homeostasis, which provides new insight into the mechanism of neurological syndromes after severe pneumonia.

## Introduction

Acute Pneumonia is mostly caused by infectious pathogens (*e.g*., bacteria, viruses, fungi), leading to an inflammatory condition of the lungs. Acute pneumonia was one of major cause of death in the past several years. In addition to acute pneumonia-related respiratory failure, neurological disorders after severe acute pneumonia are also one of the common symptoms that reduce patient quality of life. A very recent example is “brain fog” occurring in “long coronavirus disease (long-COVID)”^1,2^. Some people who have been infected with SARS-CoV-2 struggle for weeks to months with concentration issues and other cognitive symptoms, even associated with changes in brain structure^3^. In fact, neurological syndromes after severe pneumonia exist in recovered patients with many respiratory diseases^4,5^. A study showed that the frequencies of most neurological manifestations did not differ after COVID-19, influenza and bacterial pneumonia^6^, indicating a potential general mechanism involved in all acute pneumonia-induced neurological disorders. In most patients and experimental models suffering from neurological syndromes after severe acute pneumonia, hyperactive astrocytic and microglial activity is observed in the brain, accompanied by axonal damage and Aβ formation^7,8^. Given the major role of astrocytes and microglia in neuroinflammation, there is growing evidence suggesting that a detrimental immune response in the brain contributes to neurological symptoms after severe pneumonia^7^. Yet, it still remains controversial whether the alteration of the brain is induced by direct inflammatory responses from the lung or the bystander effects of pneumonia. The relationship between acute pneumonia and brain activity has not been elucidated.

Bidirectional interactions in the lung-brain axis have been reported^9^. The central nervous system (CNS) extensively communicates with other systems. Autoimmune processes are not only dependent on the local conditions of the nerve tissue itself, but also under the control of the peripheral organ system^10^. Studies have shown that the brain and lung can communicate through a variety of ways, such as neuroanatomical pathway, endocrine pathway, and immune pathway etc.^11^ The lung is an organ that contains microbiota^12^. The microbial community in the lung composes of about 140 distinct families, while most of their functions in healthy individuals are still unknown^12^. In contrast, the brain should be a sterile environment (or much fewer microbes^13^) under normal conditions, as the blood-brain barrier is able to prevent bacteria from entering the brain tissue^14^. In this study, we accidentally found that neurological disorders after severe acute pneumonia are associated with the translocation of bacteria from the lung to the brain. In the lipopolysaccharide (LPS)-induced experimental severe pneumonia mouse model, we accidentally detected an emergence of bacteria in the brain tissue. Interestingly, using 16S rDNA amplification sequencing, similarities were found between the brain’s flora species and those of the lungs, indicating the bacteria in the brain may come from the lung during pneumonia. We also observed lung-blood barrier and brain-blood barrier (BBB) impairment, simultaneously, during acute pneumonia, which may explain how lung bacteria invade the brain during acute pneumonia.

The existence of bacteria in the brain tissue results in the disruption of brain homeostasis, especially for astrocytes and microglia. Platelet-derived extracellular vesicles (PEVs) could serve as a general platform for inflammatory cells targeting drug delivery, which has been demonstrated by our group in previous studies^15–18^. By loading rapamycin into PEVs (rapamycin-PEVs), we showed that intranasal delivery of rapamycin-PEVs rescued the dysfunction of astrocytes and microglia caused by severe acute pneumonia, effectively alleviated the neurological disorders after pneumonia. Our work provides new insight into the mechanism of neurological syndromes after severe acute pneumonia, which is partly attributed to the translocation of bacteria from the lung to the brain during pneumonia.

## Results

### Severe acute pneumonia leads to neurological disorders in the brain

To avoid interference with exogenous pathogens, LPS-induced acute pneumonia was established to study neurological disorders in mice (**Figure 1A**). It was found that after LPS attack (4 mg kg^-1^), the mice experienced momentary weight loss, but quickly regained. The body weight of the treated mice was similar to that of untreated mice at day 30 after LPS attack (**Figure 1B**). The blood oxygen saturation and cytokine levels within the lungs such as IL-1β, IL-6, and TNF-α were all gradually recovered, with no difference from the untreated mice after 30 days (**Figure 1C-F**), indicating that the function of lungs in treated mice can be recovered from LPS-induced pneumonia under normal conditions. To test whether these mice suffer from a neurological disorder, the behavior of the mice recovered from pneumonia was tested. In the open field experiment, the recovered mice were found to be more huddled in the corner and immobile compared with untreated mice, with reduced total activity path, central activity path and central activity time as well as anxiety-like behavior (**Figure 1G-K**). In the new object recognition experiment, the discrimination index and preference index for new objects were significantly lower in mice that recovered from pneumonia, indicating impaired cognitive ability and short-term memory (**Figure 1L-N**). In the Morris water maze experiment, mice that recovered from pneumonia took much longer time to reach the station (**Figure 1O-P**), indicating a decrease in spatial memory ability. In addition, we also observed similar behavior abnormal in the mice after 60 days LPS treatment, indicating a long-term of the neurological disorder (**Figure S1**). Furthermore, immunofluorescence of Aβ protein in microglia and astrocytes in the brains of the recovered mice confirmed an altered function of these cells (**Figure 1Q-T**). A decrease in the length and number of microglia branches indicated that microglia were in an activated state (**Figure 1U-V**). It is noteworthy that inflammatory markers in the brain tissue after pneumonia declined, however, they were still higher than baseline (**Figure 1W-Y**). These data showed that the defect in brain function is a common feature in mice recovered from severe pneumonia.

**Figure 1.**
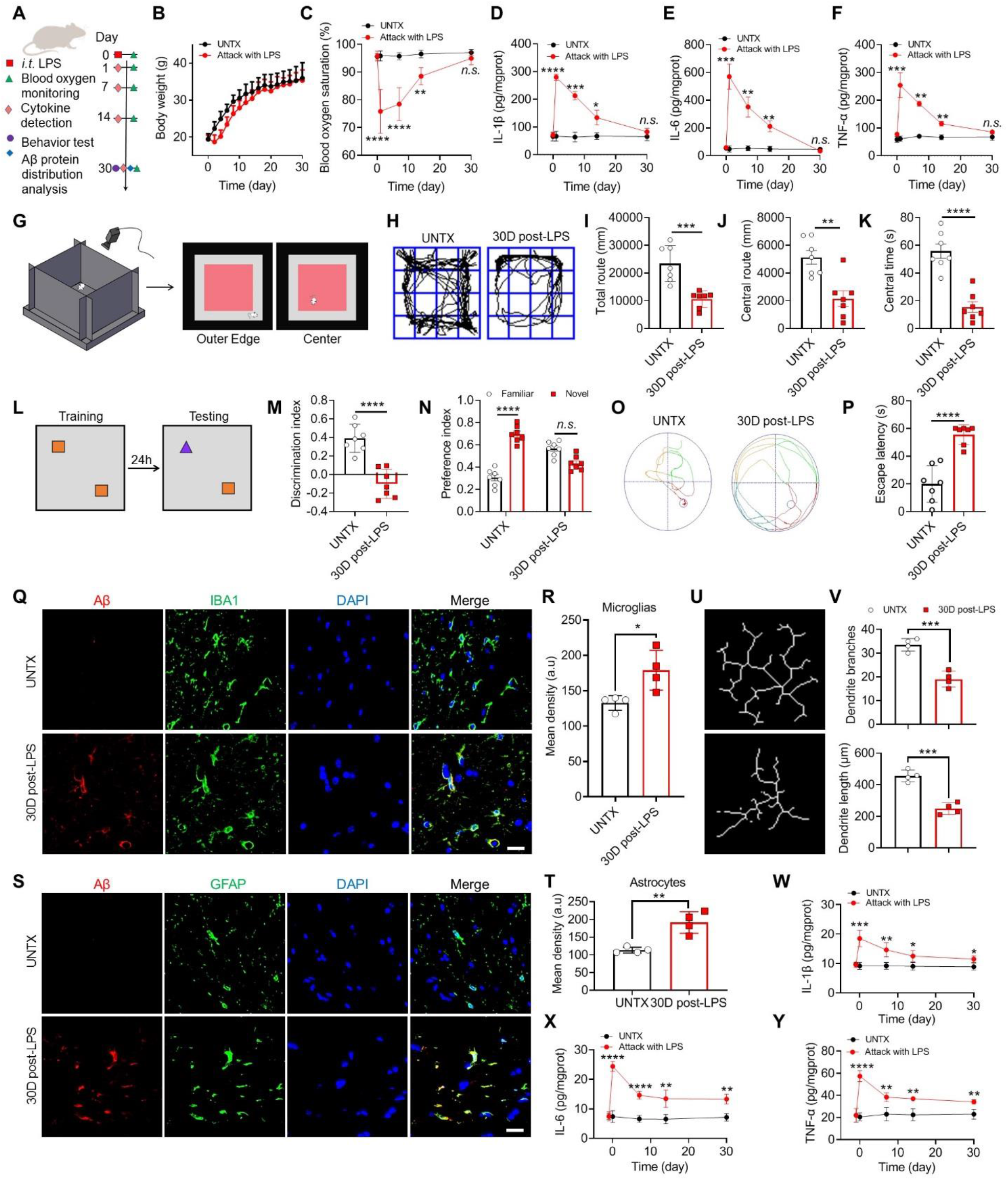
Severe acute pneumonia leads to neurological disorders in the brain. (**A**) Schematic of the experimental timeline. Severe acute pneumonia mouse model was generated by intratracheal (i.t) administration of LPS (4 mg/kg). Untreated (UNTX) and LPS-treated mice were then monitored for blood oxygen and tested for behavior, and samples were collected for molecular pathological assessment. (**B**) Body weight of untreated and LPS-treated mice. (**C**) Blood oxygen saturation of untreated and LPS-treated mice. (**D-F**) Inflammatory factors including IL-1β (**D**), IL-6 (**E**), and TNF-α (**F**) of lung tissue homogenate after LPS attack. (**G**) Schematic illustration of the open field test. (**H**) The representative paths of mice in the open field test. (**I**) The total route for mice in the open field test. (**J**) The central route for mice in the open field test. (**K**) The central time for mice in the open field test. (**L**) Schematic illustration of the novel object recognition test. (**M**) Discrimination index in the novel object recognition. (**N**) Preference index in the novel object recognition. (**O**) The representative plot of Morris water maze images and (**P**) quantification of escape latency. (**Q**) Confocal microscopy images of Aβ protein in microglia. Red: Aβ, green: IBA1, blue: DAPI. Scale bar, 20 μm. (**R**) Mean fluorescence quantification of Aβ protein in microglia. (**S**) Plot of microglia branches and (**T**) corresponding quantitative analysis. (**U**) Confocal microscopy images of Aβ protein in astrocytes. Red: Aβ, green: GFAP, blue: DAPI. Scale bar, 20 μm. (**V**) Mean fluorescence quantification of Aβ protein in astrocytes. (**W-Y**) Inflammatory factors including IL-1β (**W**), IL-6 (**X**), and TNF-α (**Y**) of brain tissue homogenate after LPS attack. Data are shown as mean ± SD (n=4-7). Statistical significance was calculated by Student’s t-test (two-tailed) and one-way ANOVA using the Tukey posttest. \**P* < 0.05; \*\**P*< 0.01; \*\*\**P* < 0.001; \*\*\*\**P* < 0.0001; n.s.; nonsignificant. a. u., arbitrary units.

### Bacteria were observed in the brain during acute pneumonia

The lung tissues harbored a microbial community that may regulate brain autoimmunity^19^. We next questioned whether alterations in homeostasis of brain was induced by the influence of the lung microbiome. Brain tissue and lung tissue was collected, homogenized, and plated on Luria-Bertani (LB) plates at different time points after LPS attack (**Figure 2A**). Bacterial colonies were quantified by counting CFUs (**Figure 2B-E**). Since the brain is considered as a sterile organ, there were no colonies in the brain of mice before LPS treatment. Unexpectedly, we observed that CFUs in the brain tissue emerged and increased obviously at days 1 and 7 after LPS attack, then gradually reduced (but not eliminated) within 30 days after pneumonia (**Figure 2B-C**). While there was no significant change in the lung CFU during the whole period of pneumonia remission (**Figure 2D-E**).

**Figure 2.**
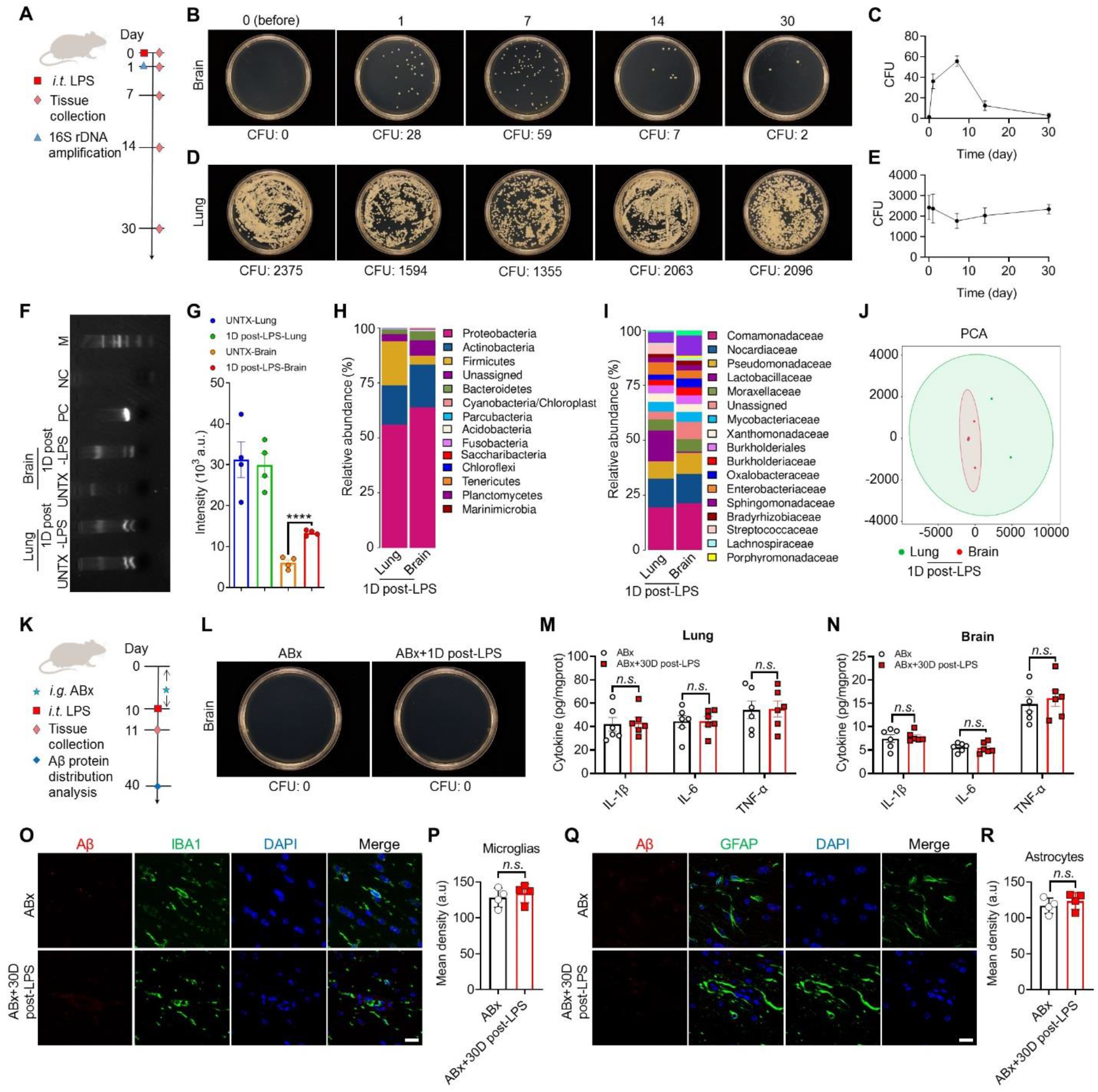
Bacteria were observed in the brain during acute pneumonia. (**A**) Schematic of the experimental timeline. Samples from untreated and LPS-treated mice were used for flora coating and 16S rDNA sequencing. (**B**) The representative plot of bacterial colony growth after 24 hours of brain tissue homogenate and (**C**) corresponding quantification results of colony forming units (CFU). (**D**) The representative plot of bacterial colony growth after 24 hours of lung tissue homogenate and (**E**) corresponding quantification results of colony forming units. (**F**) Representative agarose gel electrophoresis for DNA detection and (**G**) corresponding quantitative analysis. (**H**) Relative abundance of lung and brain bacterial inhabitants at the phylum level. (**I**) Relative abundance of lung and brain bacterial inhabitants at the family level. (**J**) Principal component analysis was performed on the lung and brain of mice 1 day after LPS attack, n=5. (**K**) Schematic of the experimental timeline. Healthy mice were treated with LPS after establishing a sterile mouse model by gavage of antibiotics, and samples were collected for molecular pathological evaluation. (**L**) The representative plot of bacterial colony growth after 24 hours of brain tissue homogenate. (**M**) Inflammatory factors expression of lung tissue homogenate. (**N**) Inflammatory factors expression of brain tissue homogenate. (**O**) Confocal microscopy images of Aβ protein in microglia. Red: Aβ, green: IBA1, blue: DAPI. Scale bar, 20 μm. (**P**) Mean fluorescence quantification of Aβ protein in microglia. (**Q**) Confocal microscopy images of Aβ protein in astrocytes. Red: Aβ, green: GFAP, blue: DAPI. Scale bar, 20 μm. (**R**) Mean fluorescence quantification of Aβ protein in astrocytes. Data are shown as mean ± SD (n=4-7). Statistical significance was calculated by Student’s t-test (two-tailed) and one-way ANOVA using the Tukey posttest. \*\*\*\**P* < 0.0001; n.s.; nonsignificant. a. u., arbitrary units.

To confirm the existence of bacteria in the brain tissue, the genomes of the lung and brain were extracted and the V3-V4 detection region (fragment in 16S rDNA gene) was selected for PCR amplification. Standard bacterial genomic DNA Mix was used as a positive control. The amplified products were subjected to agarose gel electrophoresis. The results showed that an obvious amount of bacterial genes were present in the brain genome of mice 1 day after LPS attack. In contrast, few bacterial genes were detected in the brains of untreated mice (**Figure 2F-G**). Furthermore, we evaluated the microbial composition in brain tissue and lung tissue of the mice by 16S rDNA amplicon sequencing. According to the database, the composition of the identified flora was analyzed at the level of phylum, class, order, family, genus and species (**Figure 2H-I, Figure S2**). It was found that the relative abundance between the brain and the lung flora species was very similar at the level of phylum and family classification (**Figure 2H, I**). Principal component analysis (PCA) further determined the similarity in diversity and abundance of microbiota in the brain and the lung, suggesting that the emerging bacteria in the brain may originate from the lung (**Figure 2J**). This series of experiments indicates that the brain can be invaded by bacteria that may originate from the lung after pneumonia.

The results above prompted us to explore whether alterations in the brain were caused by invading bacteria. We indiscriminately eliminated the bacteria by antibiotic cocktail treatment (**Figure S3**). Following this, the mice were administered the same dose of LPS to induce pneumonia (**Figure 2K**). We confirmed that the brain tissue of mice receiving antibiotic therapy did not have any bacteria (**Figure 2L**). After 30 days, inflammatory factors were determined in both lung and brain tissues. Notably, when the lung inflammation subsided 30 days after LPS attack, the brain inflammation of germ-free mice also recovered to baseline (**Figure 2M-N**). Immunofluorescence images of the brains revealed no significant increase in Aβ protein content in microglia and astrocytes (**Figure 2O-R**), indicating these cells were not dysfunctional. These results suggested that the alterations in the brain were associated with invaded bacteria.

### Bacteria translocated from the lung to the brain during acute pneumonia is associated with increased lung and brain permeability

We next sought to determine the mechanism by which bacteria translocated from the lung to the brain. The above observations are indicative of the scenario in which the lung-blood barrier and the blood-brain barrier are leaky during acute pneumonia, leading to an open access route for bacteria translocation. To validate this hypothesis, we analyzed the diffusion of high-molecular-weight (4 kDa) FITC-conjugated dextran (FITC-DXT) in mice following pneumonia. FITC-DXT is a general tool to assess the integrity of the blood-brain barrier (BBB)^20^. An increased permeability of the BBB is associated with accumulation of FITC-DXT in brain. In the LPS-induced pneumonia mouse model, we found that as early as 2 hours post injection, accumulation of dextran in the brain tissue was observed (**Figure 3A, Figure S4**), which was further confirmed by confocal imaging and flow cytometry of brain cells (**Figure 3B-D**). These findings are also consistent with a previous study that reported that the most prominent sign of severe COVID is BBB impairment^21^. We further found that the extent of BBB permeability was related to the magnitude of the pneumonia response. Although 0.1 mg/kg of LPS attack caused breathing difficulties in mice (**Figure S5A**), there was no significant signal of FITC-DXT in the brain. However, at a dose higher than 1 mg/kg LPS, FITC-DXT was more obviously enriched in the brain (**Figure S5B-D**), suggesting that BBB permeability correlated with the level of pneumonia, while mild pneumonia did not induce BBB impairment in our experiment.

**Figure 3.**
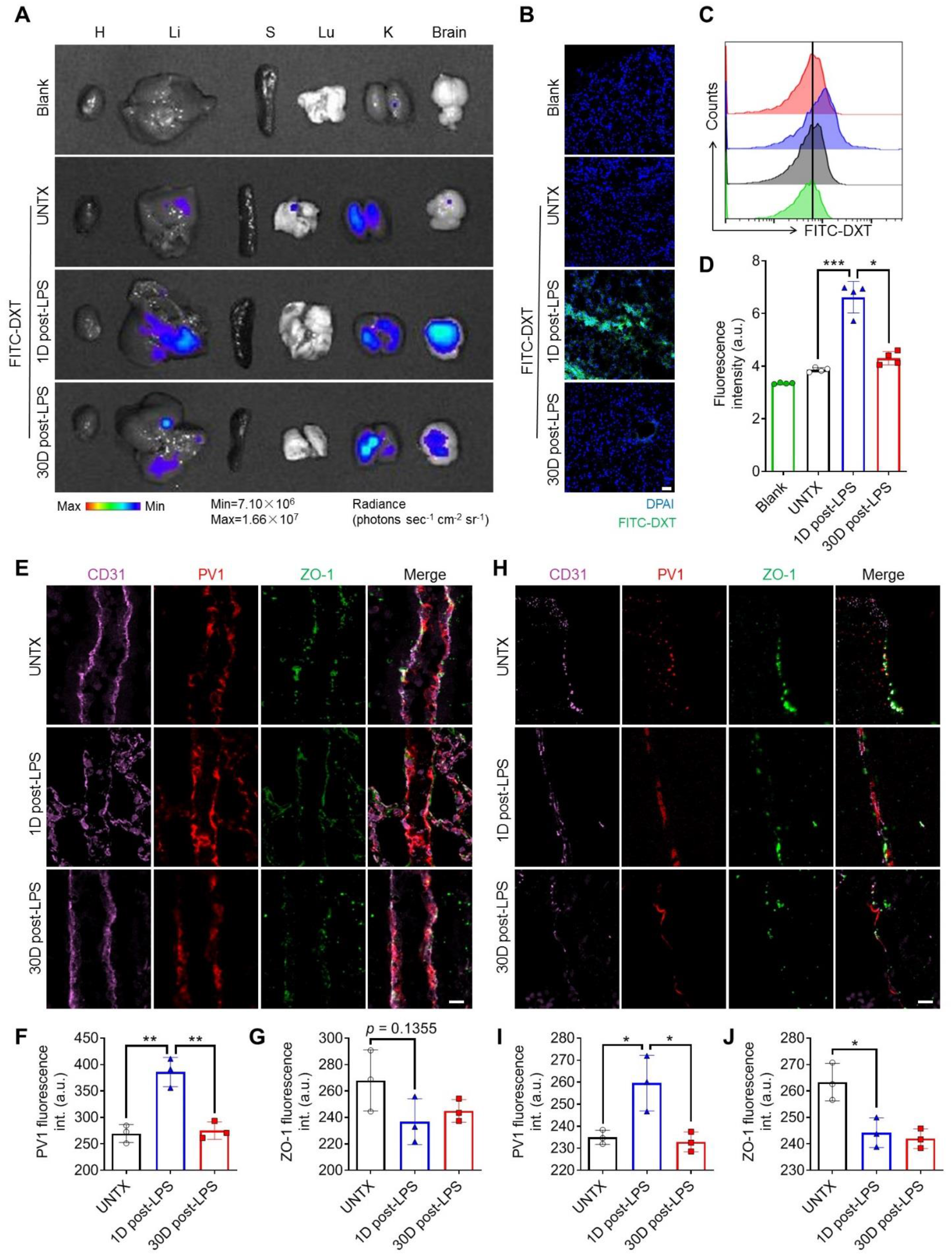
Severe acute pneumonia leads to alteration of brain and lung permeability. (**A**) *Ex vivo* imaging showed biodistribution of 4-kDa FITC–conjugated dextran. (**B**) Confocal fluorescence imaging of brain tissue slices of mice after different treatments as indicated, scale bars, 50 μm. (**C**) Representative flow cytometric analysis of 4-kDa FITC–conjugated dextran signal and (**D**) corresponding quantification results of mean fluorescence intensity (MFI) of FITC. (**E**) Representative confocal images showing PV1 (red) detection in CD31^+^ (purple) blood vessels and ZO-1 (green) in lung of mice after different treatments as indicated, scale bars, 10 μm, and (**F, G**) corresponding quantitative analysis. (**H**) Representative confocal images showing PV1 (red) detection in CD31^+^ (purple) blood vessels and ZO-1 (green) in brain of mice after different treatments as indicated, scale bars, 10 μm, and (**I**, **J**) corresponding quantitative analysis. Data are shown as mean ± SD (n=3-5). Statistical significance was calculated by Student’s t-test (two-tailed) and one-way ANOVA using the Tukey posttest. \**P* < 0.05; \*\**P*< 0.01; \*\*\**P* < 0.001; n.s.; nonsignificant. a. u., arbitrary units.

We then examined the vascular endothelial barrier and epithelial barrier in both the lung and brain one day after pneumonia. PV1 (plasmalemmal vesicle associated protein-1) is a plasma membrane vesicle-associated protein on endothelial cells, which is an indicator of inflammation-induced permeability^22^. On day 1 after LPS attack, PV1 expression increased significantly in the lung and brain (**Figure 3E-J**), indicating that the vascular barrier was disrupted. These results were consistent with the accumulation of FITC-DXT in the brain, as endothelial cells control the passage of molecules from blood to the matrix. We also observed that PV1 expression in the lung and brain returned to baseline levels on day 30 after LPS attack (**Figure 3E-J**), suggesting that the vascular barrier was closed at that time. Zonula occludens-1 (ZO-1) is involved in the tight junctions of epithelial cells^23^. We observed a decrease of ZO1 levels on day 1 as well as day 30 after LPS attack, suggesting dysfunction of tight junctions of the epithelial barrier in both the lung and brain for a long time (**Figure 3E-J**). The gut microbiota represents a significantly higher biomass than the lung microbiota. To further prove that the emerging bacteria in the brain were not from the intestinal microbiome, we also examined intestinal permeability on day 1 following pneumonia. The results showed no significant change in intestinal permeability (**Figure S6**). These results corroborate our hypothesis that both the lung-blood barrier and the blood-brain barrier are leaky during pneumonia, allowing the translocation of bacteria from the lung to the brain.

### The altered brain homeostasis induced by bacterial translocation

To further characterize the brain microenvironment of the mice suffering from neurological disorders, we performed single-cell RNA-sequencing of brain cells from the mice recovered from pneumonia at day 30. Brain cells from individual mice were barcoded before single-cell RNA sequencing using the droplet-based system of 10× Genomics. Cells were clustered based on gene expression using an unsupervised inference analysis using the Seurat v4 pipeline. The clusters were organized into 7 “meta-clusters” including astrocytes, microglia, oligodendrocytes, T cells, monocytes, neurons, and granulocytes (**Figure 4A-B, Figure S7A-C**). Although the frequency of different types of cells was not changed significantly (**Figure 4C**), the enrichment analysis of KEGG pathways of differential genes in all brain cells showed that it was related to Huntington’s disease, Parkinson’s disease, Alzheimer’s disease, retrograde nerve signaling, and GABAergic synaptic pathway (**Figure 4D**), indicating that the mice exhibited brain dysfunction and development of neurological disorders at day 30 post-pneumonia. Of note, we also observed an association with Salmonella infection pathways in KEGG analysis. In addition, as shown in the volcano map, the expression of genes (*Rhog*, *Il1a*, *Arf1*, etc.) related to the bacterial infection pathway was significantly up-regulated (**Figure 4E**). These data further confirmed our finding that bacteria existed in the brain.

**Figure 4.**
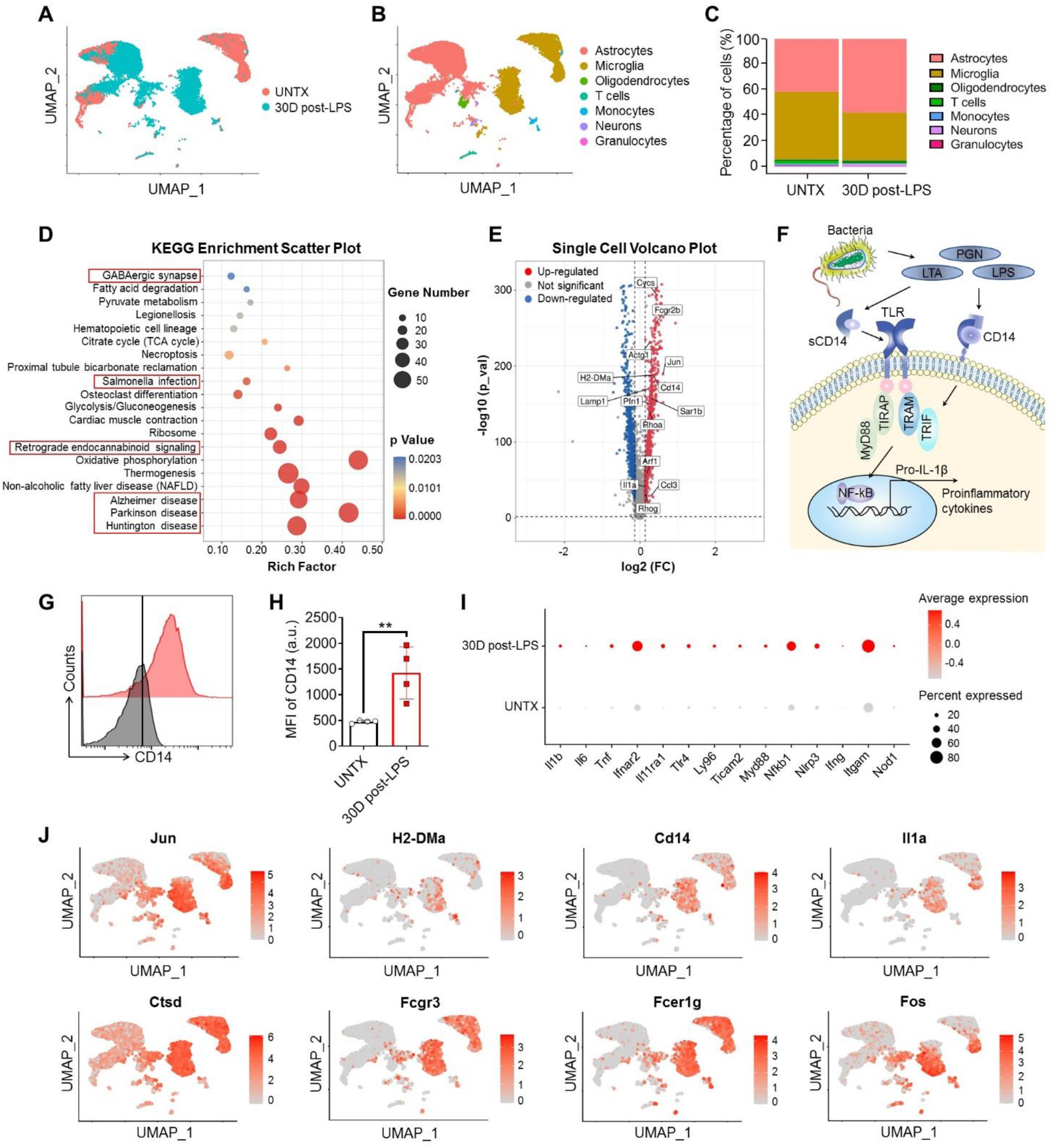
The altered brain homeostasis induced by bacterial translocation. (**A, B**) Uniform manifold approximation and projection (UMAP) showing brain cell clustering and (**C**) the percentage of each cluster. (**D**) KEGG enrichment scatter plot of brain. (**E**) Volcano plot of gene differential expression of brain. (**F**) Schematic diagram of bacterial infection. (**G**) Representative flow cytometric analysis of CD14 and (**H**) corresponding quantification results of MFI of CD14. (**I**) Dotplot of differentially expressed gene in brain. (**J**) Highlighted UMAP plot of representative genes of *Jun*, *H2-DMa*, *Cd14*, *Il1a*, *Ctsd*, *Fcgr3*, *Fcer1g*, *Fos*. Data are shown as mean ± SD (n=4). Statistical significance was calculated by Student’s t-test (two-tailed) and one-way ANOVA using the Tukey posttest. \*\**P*< 0.01.

Bacteria related pathogen-associated molecular patterns (PAMPs) can bind to pattern recognition receptors (PRRs) on various types of cells to produce pro-inflammatory factors through the NF-kB pathway^24^ (**Figure 4F**). LPS is one of representative of typical PAMP molecules derived from gram-negative bacteria. Furthermore, we verified by flow cytometry that the expression of the LPS receptor CD14 in the brains of mice was significantly increased at day 30 (**Figure 4G, H**), indicating the existence of an inflammatory immune response to bacteria in the brain tissue. In addition, the genes (*Myd88*, *Ticam2*, *Ly96*, *Tlr4*, *Nfkb1*, etc.) related to the LPS signaling pathway were all up-regulated^25^ (**Figure 4I-J**). To rule out the possibility of injected LPS that induce acute pneumonia, one day after LPS attack, the mice were given intravenous antibiotic treatment, which eliminated the bacterial flora in the brain (**Figure S8**). Based on our data, ABx-treated mice exhibited no brain inflammatory response and low CD14 expression at day 30, indicating that the LPS-related pathway was unlikely to be induced by injected LPS 30 days before.

The gene vitrification map showed that the increased expression of genes (*Jun*, *H2-DMa*, *Cd14*, *Il1a*, *Ctsd*, *Fcgr3*, *Fcer1g*, *Fos*, etc.) related to the bacterial infection pathway was mainly distributed in microglia and astrocytes which are the two primary cell types that mediate neuroinflammation (**Figure 4J, Figure S8D**). Microglial cells are the major group of macrophages in the parenchyma of the central nervous system, and are very sensitive to brain bacteria. As expected, the expression of microglial genes (*Rhog*, *Il1a*, *Arf1*, etc.) related to bacterial infection pathways were significantly up-regulated (**Figure 5A**), and KEGG pathway showed that this cluster was involved in Salmonella and Staphylococcus infections (**Figure 5B**). At the same time, microglial cells were activated as shown by a variety of markers, such as *C1qa*, *C1qb*, *C1qc* (complement pathway genes), *Ctss*, *Ctsb* (lysosome pathway genes), *Ccl3*, *Ccl4* (chemokines), *Rtp4*, *Bst2* (interferon response genes), *Lamp1*, *Lamp2*, *H2-D1*, *P2ry12*, *Hexb*, *Trem2* (microglia activation related genes)^26,27^ (**Figure 5C, Figure S9A**), and NLRP3 dominated the inflammasome pathway (**Figure S9A**). Similarly, in astrocytes, we found that the expression of genes (*Rhog*, *Il1a*, *Arf1*, etc.) associated with bacterial infection pathways was also remarkably upregulated (**Figure 5D**). The KEGG pathway analysis of astrocytes showed that this cluster is involved in Huntington’s disease, Parkinson’s disease, Alzheimer’s disease, retrograde nerve signaling, GABAergic synapses, and lysosomal pathways (**Figure 5E**), indicating that astrocyte reactivity is associated with neurological disorders. Moreover, the expression of reactivity marker genes of astrocytes, such as *Id3*, *Npc2* and *Prdx6*^28^, Alzheimer’s disease risk-related genes *Ctsb*, *Ctsd*, *Ctsl*, *S100a6*, *Itgab5* and *Vsir*^29^ and interferon stimulated genes *Gbp3*, *Gbp2*, *Irgm1*, *Iigp1*, *Igtp*, *Cxcl10* were all up-regulated in the astrocyte of brain^30–32^ (**Figure 5F, Figure S9B**). All these data suggest that the bacteria are involved in alternations of brain homeostasis, by switching both microglia and astrocytes from quiescence to activation though bacterial infection-related pathways. Meanwhile, the activation of microglia and astrocytes is linked to neurological diseases.

**Figure 5.**
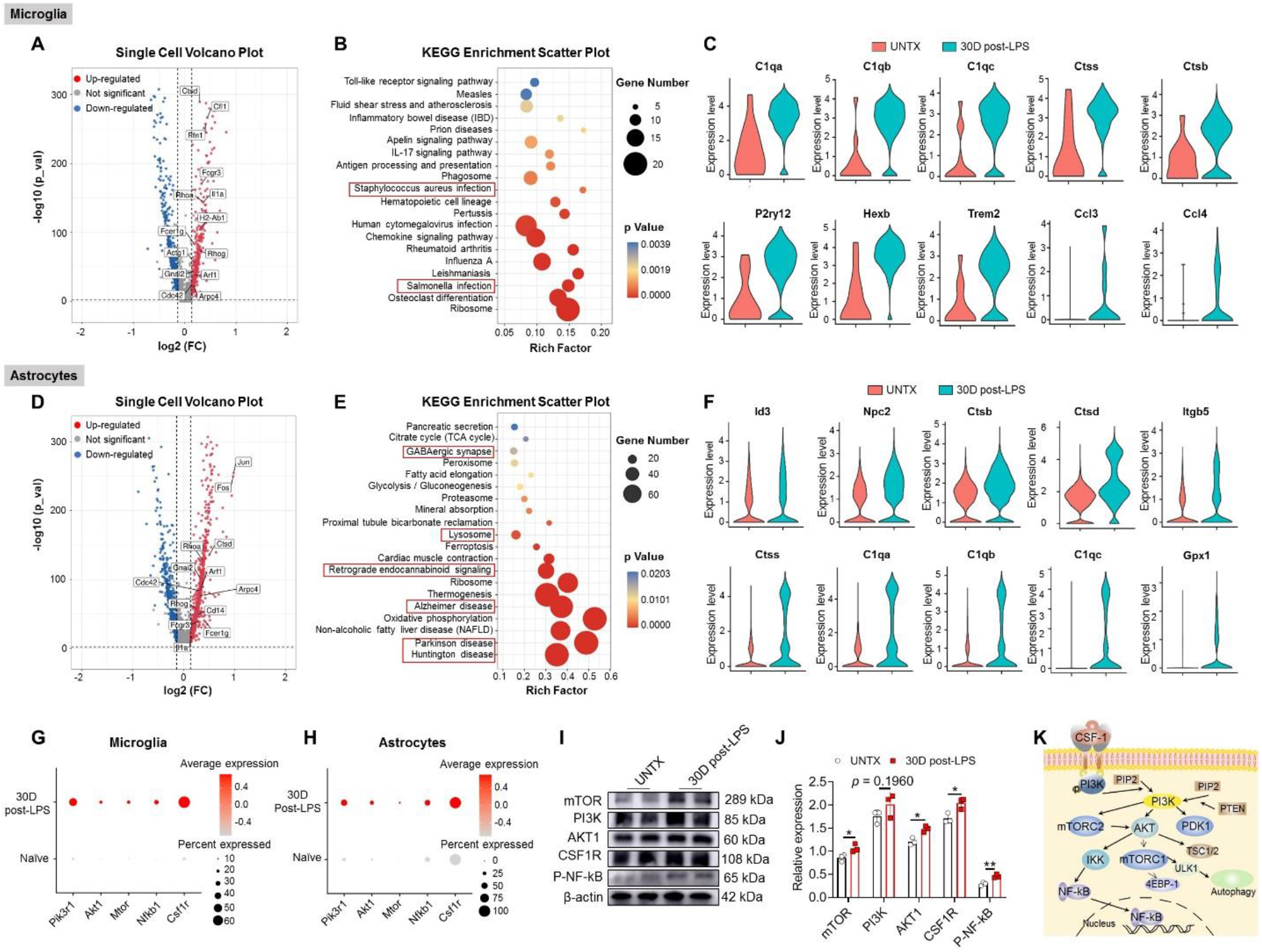
Dysfunction of astrocytes and microglia. (**A)** Volcano plot of gene differential expression of microglia. **(B**) KEGG enrichment scatter plot of microglia. **(C**) Violin plots of representative differential gene expression in microglial cluster. (**D**) KEGG enrichment scatter plot of astrocytes. (**E**) Volcano plot of gene differential expression of astrocytes. (**F**) Violin plots of representative differential gene expression in astrocytes cluster. (**G**) Dotplot of differentially expressed gene in microglia. (**H**) Dotplot of differentially expressed gene in astrocytes. (**I**) Schematic diagram of the mTOR signaling pathway. (**J**) Western blot analysis of the expression of various types of proteins in brain cells after various treatments as indicated and (**K**) the relative expression of proteins compared to the untreated group. Data are shown as mean ± SD (n=3). Statistical significance was calculated by Student’s t-test (two-tailed) and one-way ANOVA using the Tukey posttest. \**P* < 0.05; \*\**P*< 0.01.

In particular, we found upregulation of PI3K-AKT-mTOR signaling in both microglial (**Figure 5G**) and astrocytic (**Figure 5H**) clusters, which was further validated by Western Blotting approach (**Figure 5I-J**). Numerous studies have shown that the PI3K-AKT-mTOR pathway is a key signaling pathway utilized by microglia and astrocytes to respond to extracellular stimuli including bacteria. This pathway is considered central for maintaining brain homeostasis while abnormal PI3K-AKT-mTOR signaling linked to various neurological diseases, such as Alzheimer’s disease, Parkinson’s disease, and Huntington’s disease. Given the central role of PI3K-AKT-mTOR signaling in neurological homeostasis (**Figure 5K**), it may be a therapeutic target in pneumonia-induced neurological disorders.

### Rapamycin rescues brain homeostasis and neurological disorders

Rapamycin is used extensively for selective mTOR inhibition. Studies have shown that rapamycin reduces inflammation and inhibits the pathological processes of neurodegenerative disorders by inhibiting mTOR^33,34^. Therefore, we hypothesized that treatment with rapamycin may rescue the pneumonia-induced neurological disorders (**Figure 6A-B**). As the rapamycin we used here was relatively insoluble and unstable (**Figure 6A**), its distribution in the brain was quite limited by intravenous injection in our preliminary data (data not shown). In our previous studies, we demonstrated that platelet-derived extracellular vesicles (PEVs) could serve as a universal platform to selectively target various inflammatory cells and tissues^15–18^ (**Figure 6C**). To effectively deliver rapamycin to the brain, PEVs were used as carriers for direct nasal delivery of rapamycin, which is an effective and reliable way to bypass the blood-brain barrier and deliver drugs into the central nervous system^35^.

**Figure 6.**
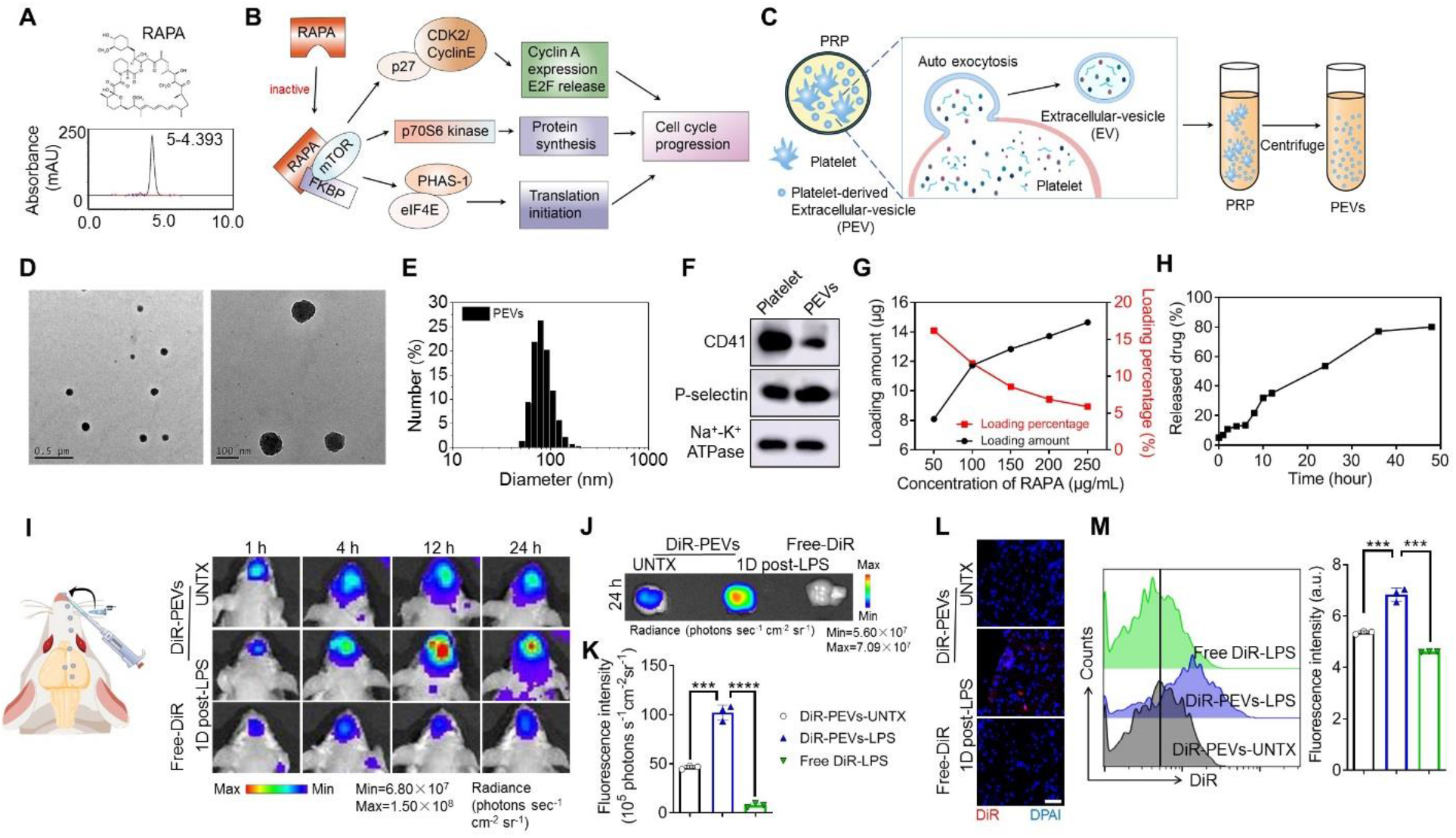
Characterization and brain accumulation of rapamycin-PEVs. (**A)** Chemical structure and HPLC characteristic peak of rapamycin. **(B**) Rapamycin mechanism diagram. **(C**) Scheme of the preparation of PEVs. (**D**) Morphology of PEVs by TEM. (**E**) PEVs size distribution measured by DLS. (**F**) Western blot results of platelet lysate and PEVs. (**G**) Drug loading amount and efficacy of RAPA to the PEVs. (**I**) Schematic diagram of intranasal administration and *in vivo* fluorescence imaging of the mice after intranasal administration DiR labelled PEVs. (**J**) *Ex vivo* fluorescence imaging of brain and (**K**) corresponding quantitative analysis. (**L**) Confocal fluorescence imaging of brain tissue slices of mice after different treatments as indicated, scale bars, 50 μm. (**M**) Representative flow cytometric analysis of DiR-PEVs and corresponding quantification results of MFI of DiR. Data are shown as mean ± SD (n=3). RAPA: rapamycin. Statistical significance was calculated by Student’s t-test (two-tailed) and one-way ANOVA using the Tukey posttest. \*\*\**P* < 0.001; \*\*\*\**P* < 0.0001.

Transmission electron microscopy (TEM) and dynamic light scattering (DLS) analysis showed that PEVs had a round shape with a diameter of approximately 80–150 nm (**Figure 6D-E**). PEVs retained adhesion molecules from platelets, including CD41 and P-selectin (**Figure 6F, Figure S10A**), which can bind to activated microglia and astrocytes. According to our established protocol^15–18^, rapamycin can be loaded onto PEVs by hydrophobic interactions (**Figure S10B**). The loading did not change the PEVs significantly (**Figure S10C, D**). At a rapamycin concentration of 100 μg mL^-1^, the drug loading rate (loading/adding rapamycin) was approximately 11.73% (**Figure 6G**). Meanwhile, 80.01% of rapamycin was released from PEVs within 48 h, showing a sustained release profile (**Figure 6H**).

We next examined brain accumulation of the rapamycin-PEVs *via* intranasal (IN) administration (**Figure 6I**). In our experiment, DiR-labelled rapamycin-PEVs were IN administrated to the mice with pneumonia on day 1. Naïve mice were used as controls. The mice were imaged at the experimentally designed time points using a *in vivo* near infrared fluorescence system. As expected, a remarkable accumulation of DiR-rapamycin-PEVs was observed in the brains of pneumonia mice, which was more significant than that in normal mice (**Figure 6I**). This can be explained by a targeting effect of PEVs to activated microglia and astrocytes. In addition, the PEVs in the brain reached the maximum enrichment at 12 h, and the retention time of PEVs was significantly longer than that of controls (**Figure 6I**). *Ex vivo* NIRF imaging also revealed substantial accumulation of PEV in the brains of mice with pneumonia (**Figure 6J-K**). These findings are supported by confocal imaging (**Figure 6L**) and flow analysis, which showed a higher number of DiR+ cells in the brain after intranasal administration (**Figure 6M**). Together, these data suggested that rapamycin can be effectively delivered into brain tissue with the help of PEVs.

Next, we assessed the therapeutic efficacy of rapamycin-PEVs on mice suffering from pneumonia-induced neurological disorders (**Figure 7A**). One day after LPS attack, mice received intranasal administration of rapamycin-PEVs once every two days for a total of four times. Naïve mice, untreated mice, and mice treated with PEVs or rapamycin alone were used as controls. Behavioral tests were performed on day 30. The results of the open field experiment showed that rapamycin-PEVs effectively alleviated the reduced motor activity induced by pneumonia-induced neurological disorder in untreated mice (**Figure 7B-E**). For the water maze, intranasal administration of rapamycin-PEVs significantly rescued the probability of mice entering the target quadrant and reduced the time required to reach the target platform compared to the untreated mice, while there was no difference in swimming speed (**Figure 7F-I**). In addition, mice receiving rapamycin-PEVs significantly mitigated the reduction in the new object preference index after pneumonia (**Figure 7J-K**). Free rapamycin or PEVs alone had inferior or limited therapeutic effects (**Figure 7B-K**). These results suggest that intranasal delivery of rapamycin-PEVs can significantly rescue pneumonia-induced behavioral disorders.

**Figure 7.**
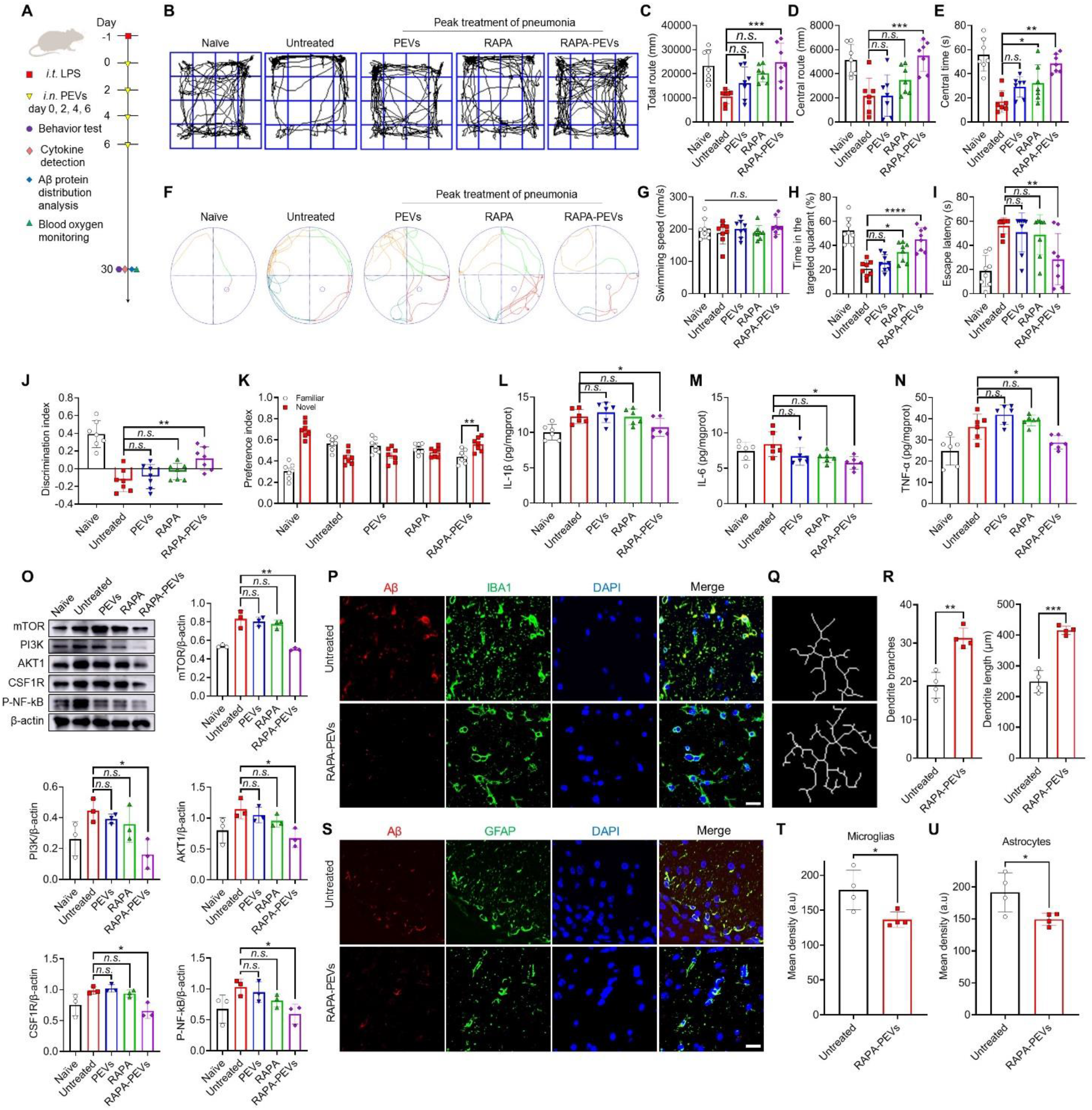
Therapeutic effect of rapamycin-PEVs. (**A**) Schematic of the experimental timeline. One day after LPS attack, mice received intranasal administration of rapamycin-PEVs once every two days for a total of four times. Naïve mice, untreated mice, and mice treated with PEV or rapamycin alone were used as controls. Behavioral testing and sample collection were conducted on day 30. (**B**) The representative paths of mice in the open field test and quantitative analysis of (**C**) the total route, (**D**) the central route and (**E**) the central time. (**F**) The representative plot of Morris water maze images and quantification of (**G**) swimming speed, (**H**) the ratio of the time in the target quadrant to the total time, and (**I**) escape latency. (**J**) Discrimination index in the novel object recognition. (**K**) Preference index in the novel object recognition. (**L-N**) Inflammatory factors including IL-1β (**L**), IL-6 (**M**), and TNF-α (**N**) of brain tissue homogenate. (**O**) Western blot analysis of the expression of various types of proteins in brain cells after various treatments as indicated. (**P**) Confocal microscopy images of Aβ protein in microglia. Red: Aβ, green: IBA1, blue: DAPI. Scale bar, 20 μm. (**Q**) Plot of microglial branches and (**R**) corresponding quantitative analysis. (**S**) Confocal microscopy images of Aβ protein in astrocytes. Red: Aβ, green: GFAP, blue: DAPI. Scale bar, 20 μm. (**T, U**) Mean fluorescence quantification of Aβ protein in microglia (**T**) and astrocytes (**U**). Data are shown as mean ± SD (n=4-7). RAPA: rapamycin. Statistical significance was calculated by Student’s t-test (two-tailed) and one-way ANOVA using the Tukey posttest. \**P* < 0.05; \*\**P*< 0.01; \*\*\**P* < 0.001; n.s.; nonsignificant. a. u., arbitrary units.

In addition, rapamycin-PEVs administration also recovered the levels of the pro-inflammatory cytokines IL-1β, IL-6, and TNF-α in the brain homogenate at day 30 (**Figure 7L-N**). Moreover, the expression of mTOR, the target protein of rapamycin was decreased, and the expression of PI3K, AKT1, CSF1R, and P-NF-kB was down-regulated (**Figure 7O**), indicating that rapamycin-PEVs treatment could effectively relieve the microglial and astrocytic dysfunction in the brain following the pneumonia. In addition, the expression of Aβ protein in the mouse brain decreased after rapamycin-PEVs treatment, and the increase of microglial branch length and number indicated that microglia restored homeostasis (**Figure 7P-U**). Rapamycin-PEVs treatment also showed good safety in treated mice (**Figure S11**). These data suggest that intranasal delivery of rapamycin-PEVs showed impressive therapeutic efficiency in rescuing the brain homeostatic processes disrupted by bacterial translocation.

## Discussion

Brain-related abnormalities are frequently reported by patients who have recovered from pneumonia. For example, the inability to concentrate is the major problem of people recovering from COVID-19 in the United States and Europe^36^. In fact, neurological disorders also exist in recovered patients with other respiratory diseases, such as influenza^37^ and bacterial pneumonia^5^. In this paper, we accidentally found that neurological disorders after severe pneumonia are associated with the translocation of bacteria from the lung to the brain. The lung is frequently exposed to air and contains a pulmonary microbial community, while its function is still controversial. Changes in the pulmonary microbial community seem to play a role in progression of pulmonary disorders^12^. In contrast, the brain should be a sterile environment under normal conditions, as the blood-brain barrier is able to prevent bacteria from entering the brain tissue. However, in acute pneumonia condition, we detected bacteria in brain tissue. We observed bacterial growth in the LB plates containing mouse brain tissue homogenizing fluid following days of pneumonia. The peak of the bacterial amount in the brain was approximately 7 days post pneumonia. The existence of bacteria in the brain tissue was further confirmed by 16S rDNA gene detection and analysis. To rule out the possibility of exogenous bacterial contamination during surgery or operation, single-cell RNA sequences of brain cells clearly indicated that genes related to the bacterial infection pathway were significantly up-regulated even 30 days later. More interestingly, from the 16S rDNA gene analysis and principal component analysis, we found a similarity in the diversity and abundance of microbiota in the brain and lung, which is indicative of the scenario that the emerging bacteria in the brain originate from the lung.

In previous studies, bacteria or their products severed as direct mediators of the brain-lung axis. Cryptococcus is a fungal pathogen that causes disease in humans and enters the body primarily through inhalation. In some cases, it progresses to pneumonia followed by the spread of the infection to the central nervous system, resulting in meningoencephalitis^38^. Other evidence suggests that the lung microbiome contributes to brain-related diseases. Chronic obstructive pulmonary disease (COPD) alters the respiratory microbiota to increase the risk of Parkinson’s disease (PD) and Alzheimer’s disease (AD)^39^. In addition, studies have confirmed that similar microbiomes were detected in both bronchoalveolar lavage fluid (BAL) and blood in LPS-induced pneumonia, suggesting the migration of bacteria from the lung to the blood circulation^40,41^. In our study, we further validated that both the lung-blood barrier and the blood-brain barrier are leaky during pneumonia, allowing the translocation of bacteria from the lung to the brain *via* the bloodstream.

Furthermore, the level of BBB leakage is highly correlated with the severity of pneumonia; the more severe the pneumonia, the more leaking the BBB. This phenomenon may explain the data of many previous studies. For example, a study of the medical records of US veterans analyzed various health burdens in the six months after COVID-19 infection, which clearly indicated that neurological abnormalities were significantly higher in severe (requiring hospitalization) patients than in mild (not requiring hospitalization) patients^42^. Another study analyzed long COVID symptoms in 2020 and 2021, which once again showed that the proportion of sequelae was related to the severity of pneumonia at the time of infection^36^.

The existence of bacteria in brain tissue results in the disruption of brain homeostasis, especially for astrocytes and microglia which are the two primary cell types that mediate neuroinflammation^43^. According to an automatedsingle cell RNA sequencing study, dysregulated astrocytic and microglial signatures were displayed in COVID-19 brains^44^. Activated microglia and astrocytes are sensitive indications in response to bacterial infection, by producing cytokines, chemokines and reactive oxygen species that are beneficial in killing bacteria^45,46^. However, by-product of this local inflammation results in short-term damage to the CNS and neurological diseases^47,48^. Even though the bacteria in the brain can be eliminated completely by the activated immune system, the switching of microglia and astrocytes from activation to quiescence takes more time. A study has shown that patients with post-COVID-19 brain fog recover completely over the course of 6 to 9 months^49^, indicating that these neurological disorders can be recovered after a period of time, which is consistent with our findings.

To prevent potential damage to the normal CNS and speed up the process of microglia and astrocytes from activation to quiescence, we used rapamycin as a therapeutic agent for treatment. We found the upregulation of PI3K-AKT-mTOR signaling in both microglia and astrocyte clusters from single-cell RNA sequencing data, while PI3K-AKT-mTOR is a key signaling pathway for maintaining brain homeostasis. These data suggest that mTOR may be a therapeutic target in pneumonia-induced brain dysfunction. Rapamycin as a selective mTOR inhibitor has been demonstrated to treat pathological processes of neurodegenerative changes previously^33,34^. In addition, we used PEVs as a drug delivery platform for effectively delivering rapamycin into brain tissue, thus reducing the dosage and potential side-effects of rapamycin to the whole body. Behavioral impairments in mice were alleviated by intranasal administration of rapamycin-PEVs, thereby providing an effective low-cost approach to reduce potential damage in the brain during pneumonia. Other clinically approved anti-inflammatory agents can be examined in the future.

This study has a few limitations that warrant discussion. First, we only used LPS to establish a pneumonia model; however, whether virus-induced pneumonia has a similar effect needs further examination. Second, although some viruses, such as SARS-CoV-2, have rarely been detected in the cerebrospinal fluid of patients with neurological symptoms^50,51^, we cannot rule out the possibility that the other exogenous pathogenes translocate into brain^52^. In addition, microbial nutrients or their derivatives, such as short-chain fatty acids or bacterial outer membrane vesicles (OMVs), can also influence the activation of astrocytes and microglia, which should be investigated in the feature.

In conclusion, our study found that neurological disorders after severe pneumonia were associated with the translocation of bacteria from the lung to the brain. We observed the existence of bacteria in the brain tissue, which might result from the leakage of both the lung-blood barrier and the blood-brain barrier during pneumonia. Using single-cell RNA sequencing technology, we identified PI3K-AKT-mTOR signaling pathway disruption in both microglia and astrocyte clusters. Administration of rapamycin-PEVs could speed up the recovery of brain homeostasis, and alleviate behavioral impairments in mice suffering from severe pneumonia-induced brain symptoms. Our work provides new insights into the mechanism of neurological syndromes after severe pneumonia, which is partly attributed to the translocation of bacteria from the lung to the brain.

## Materials and Methods

### Materials

Rapamycin and antibodies applied in this study are shown in **Supplementary Table 1**.

### Animals

Female KM mice aged from 4 weeks were purchased from Nanjing Peng Sheng Biological Technology Co., Ltd. There was an at least 7-day gap between the time of purchasing mice and our experiment on them to ensure that they were accustomed to the conditions of the laboratory. The mice were housed in a vivarium maintained at 20 ± 2°C, 55% humidity, with a 12-h light–dark cycle and free access to food and water. The housing group was five at maximum for mice in each group. All animal tests were conducted with the approval of Soochow University Laboratory Animal Center and the Institutional Review Committee, in accordance with relevant ethical and moral standards (No. SUDA20200512A01).

We used the ARRIVE reporting guidelines. The aim of this study was to investigate the relationship between neurological disorders following severe pneumonia and the pulmonary microbial community. Animal experiments shall be approved and supervised by the Animal Welfare Review Committee after the purpose, method and ethics of the experiments are clarified. Mice were randomly divided into groups. No animals were excluded from the study, and the researchers conducted the experiments independently and evaluated the results. The mice were euthanized at the end of the experiment or when they had health problems. All experiments were repeated at least 3 times.

### Animal Model Induction and Treatment

For LPS treatment, healthy mice were anesthetized with isoflurane. We placed each mouse in an air-numbed chamber and adjusted the oxygen flowmeter to between 0.6 to 1.2 L/min. Once fully anesthetized, the mice were fixed in the supine position. The mouth of the mice was opened, the tongue was picked out with forceps and placed in the lateral position, and the exposed tracheal hole was observed under spotlight. LPS (4 mg/kg, Biosharp) 50 μL was injected into the trachea through a syringe. LPS challenged mice and untreated healthy mice as controls were weighed at the same time and monitored every two days until 30 days after modelling. In addition, blood oxygen saturation was measured by a MouseOx^®^ (STARR) oxygen detector at days −1, 0, 7, 14 and 30 of LPS attack. The mice were included in the study if the blood oxygen saturation dropped remarkably, and these mice were randomly divided into four groups, treated with intranasal/pulmonary delivery of rapamycin (1 mg/kg, Energy chemical) or rapamycin–PEVs (equal to 1 mg/kg of rapamycin), respectively. Some healthy mice of the same age without any treatment were utilized as controls.

### Antibiotic Cocktail Therapy

As previously described, 12 mice were gavaged with sterilized drinking water supplemented with the following broad-spectrum antibiotic cocktails: ampicillin (1 mg/mL, Aladdin), gentamicin (1 mg/mL, Aladdin), metronidazole (1 mg/mL, Energy Chemical), and vancomycin (0.5 mg/mL, Aladdin) continuously administered for 10 days to prepare germ-free mice. In addition, 6 sterile mice were selected for acute pneumonia modelling, and appropriate experiments were conducted 1 day and 30 days after modelling. The remaining 6 sterile mice were used as controls. In addition, 8 acute pneumonia mouse models were established separately. 1 day after the modelling, 4 acute pneumonia mice were randomly selected to receive tail vein antibiotic treatment, and then appropriate experiments were conducted 1 day and 30 days after treatment. The remaining 4 acute pneumonia mice were used as controls.

### Cytokine Detection

After the tissue samples were rinsed with pre-cooled PBS, samples with the same weight were weighed and ground in a tissue grinder to make a 10% tissue homogenate. The prepared homogenate was centrifuged at 6000 × g for 10 min. The supernatant of each homogenate sample was collected, and the total protein content of each tissue homogenate sample was determined by the BCA method. After that, the reaction was performed on the pre-coated enzyme-labelled plate, followed by washing, enzyme-labelled antibody coupling, substrate reaction, and termination of the reaction operation. Finally, the multifunctional enzyme-labelled instrument was used for detection at the wavelengths of 450 nm and 570 nm.

### Behavioral Tests

Each group of 7 mice underwent the open field test, new object recognition test and Morris water maze test in turn. Before the experiment, the mice were acclimated in the test room. The experiment will be conducted on consecutive days, and the same batch of mice will be used for testing. The tests were run in the same order as on day one. In addition, behavioral tests themselves affect the brain, so we performed brain tissue analysis on mice that did not undergo behavioral tests.

### Open Field Test

The experimental device was a three-dimensional blank rectangular open field box (60 cm by 60 cm by 60 cm). The mice were quickly placed in the central area of the experimental box, and the tracking software was opened to automatically record the behavior of the mice in the box for 10 min. The open field box was wiped between each group of tests to remove the residual odor of mice. Tracking software was used to collect the following data: central movement time, central movement distance, total movement distance and movement speed.

### Novel Object Recognition Test

The day before training, the mice were exposed to an open field box for 10 min without an object to form a habit. During the training phase, in the presence of two identical objects, the mice were placed in an open field box and allowed to explore the object for 5 min. 24 h later, one of the objects was replaced with a new one, and the mice were again placed in the open field and allowed to explore for 5 min. The discrimination index and preference index were used to assess novel object recognition, and this index illustrates the differences in exploration time. Tracking software was used to collect behavioral data of mice, and statistical time was used to evaluate the mice’s preferred behavior. The discrimination index was calculated as the time spent exploring the novel object minus the time spent exploring the familiar object, divided by the total exploration time. The preference index was calculated as the proportion of total time spent exploring new or old objects. [Discrimination index = T_novel_ – T_familiar_/(T_novel_ + T_familiar_), Preference index = T_novel_ or T_familiar_/T_novel_ + T_familiar_)]. All discrimination index values fall between −1 and +1, and preference index values fall between 0 and 1.

### Morris Water Maze Test

The water maze was located in a separate laboratory to ensure that the experiment was quiet and free from influence. The pool was divided into four quadrants, and different symbols (circle, pentagram, triangle, and square) were affixed to the walls of each quadrant to provide additional spatial clues to the maze. The water temperature was maintained at 23 ± 1°C, and all water maze experiments were conducted daily at the same time points. Before the experiment, the mice were conditioned to a water maze room for 2 h. Each mouse was trained to find a hidden target platform for 5 days with two tests per day, with 4 h intertrial interval. The mice were placed gently in either of the pool quadrants, protected from human influence and facing away from the pool wall. During training, if the platform was found by the mouse for more than 60 seconds, the mice were guided onto a platform and stayed there for 10 seconds. 24 h after the training, the platform was removed and the 60-s exploratory test began. The mice were placed in water facing the quadrant opposite the target quadrant. The time spent in the target quadrant and the time taken to reach the platform location were recorded as indicators of spatial memory.

### Immunofluorescence Analysis

The tissue was fixed with 4% paraformaldehyde and then buried overnight using optimal cutting service temperature compound (OCT). The embedded organ was then cut into 4 μm slices using a cryoslicer. After being sealed with 10% fetal bovine serum (FBS) for 20 min, the slices were incubated with rabbit anti-Aβ, anti-IBA1, anti-GFAP, anti-CD31, anti-PV1 and anti-ZO1 (1:500, applicable to each of these antibodies, Serviceio) at 4°C overnight. After being placed at room temperature for 30 min on the second day, the slices were incubated with a fluorescent goat anti-rabbit secondary antibody (1:500, Serviceio) for 2 h, and incubated with 4,6-diamidino-2-phenylindole (DAPI, 1 μg/mL, Beyotime) for 10 min. Images were obtained by confocal microscopy, and the image data were processed by Image J software package. In addition, the behavior test itself affects the brain, so we performed immunofluorescence analysis of brain tissue in mice that did not undergo the behavior test.

### Tissue Sample Plate Coating

Mouse tissue samples were weighed on a sterile super clean table and ground into a 5% tissue homogenate liquid. The sample was diluted with sterilized water. A certain amount of sample liquid was absorbed with a sterile pipette and coated evenly on a plate containing LB medium (Beyotime) prepared in advance after high temperature sterilization. All reagents and consumables were sterilized before use, and the entire operation was carried out on an ultra-clean platform to ensure no bacterial contamination. LB medium was inverted in an incubator for 24 h, and quantitative colony forming units were analyzed using colony counter.

### 16S rDNA Amplicon Sequencing

16S rDNA amplicon sequencing was performed by Genesky Biotechnologies Inc., Shanghai, 201315 (China). In short, total genomic DNA was extracted using the FastDNA^®^ SPIN Kit for Soil (MP Biomedicals, Santa Ana, CA) according to the manufacturer’s instructions. The integrity and quality of genomic DNA were determined by agarose gel electrophoresis, and the concentration and purity of genomic DNA were determined by Nanodrop 2000 and Qubit3.0 Spectrophotometer. The V3-V4 hypervariable regions of the 16S rDNA gene were amplified with the primers 341F (5’-CCTACGGGNGGCWGCAG-3’) and 805R (5’-GACTACHVGGGTATCTAATCC-3’) and then sequenced using an Illumina NovaSeq 6000 sequencer.

### Brain Permeability Assay to 4 kDa Dextran Molecule

According to the experimental design, 4 mice were in each group, including untreated mice, mice treated with 0.1 mg/kg, 1 mg/kg, 4 mg/kg LPS after 1 day and 4 mg/kg LPS after 30 days, respectively. 500 ug of 4 kDa FITC dextran (MCE) was injected into the tail vein of mice. 12 hours after the injection, the major organs of mice were collected for NIRF imaging. Then, the brain tissues were sectioned by OCT and observed under a confocal microscope with DAPI staining. Fluorescence histograms of brain cells were recorded using BD FACSCalibur flow cytometer, and FITC fluorescence was detected using Flowjo_V10 software based on 10,000 gated events.

### Single Cell RNA Sequencing of Brain

Single-cell RNA sequence brains were obtained from untreated or 30D post-LPS-treated mice. The tissue was digested with enzymes and processed into a single cell suspension. The oligo dT-based complementary DNA database is a droplet-partioning barcode using the Chromium Single Cell Controller (10-fold genomics) system in the NCI-CCR single cell analysis tool. Carry out the removal of dead cells, adjust the cells to the required concentration, and approximately 8000 cells were detected on the machine. Sequencing was performed on a Nova Seq (Illumina) at the NCI-CCR sequencing facility. Bioinformatic analysis was performed using the OmicStudio tools at https://www.omicstudio.cn/tool.

### Western Blotting

After the brain tissue of each experimental group was obtained, a mixed solution of radio immunoprecipitation assay (RIPA) lysate and phenylmethanesulfonylfluoride (PMSF) (100:1) was added. After lysis in an ice bath, the protein supernatant was obtained by centrifugation at 12000 rpm, and the protein was quantified by a BCA protein quantification kit. The denatured protein was mixed with 5× load buffer and isolated by 12.5% SDS-PAGE gel (Epizyme) electrophoresis at 60 V for 30 min, 120 V for 90 min. After that, the protein was transferred to PVDF membrane by ice bath for 100 min and then blocked in 5% skim milk powder solution for 1 h. After that, the protein was incubated with β-actin (1:2000, Serviceio), anti-mTOR (1:1500, Serviceio), anti-PI3K (1:1500, Abclonal), AKT1 (1:1500, Serviceio), CSF1R (1:1500, Abclonal), P-NF-KB antibodies (1:1500, Abclonal) at 4□ overnight. After that, goat anti-rabbit secondary antibody (1:5000, Absin) coupled with horseradish peroxidase was incubated at room temperature for 1 hour, bands were displayed by chemiluminescence development, and data quantitative analysis was performed by Image J software package (**Figure S12**).

### Platelet-Derived Extracellular Vesicles Fabrication

Extracellular vesicles of platelet origin were collected through natural platelet activation secretion and superfast centrifugation concentration. In short, fresh blood was collected from the fundus venous plexus of healthy KM mice, then centrifuged at 100 × g for 15 min to extract supernatant platelet-rich plasma (PRP). The pelleted PRP was resuspended in PBS containing prostaglandin E1 (2 μM, Absin) and ethylene diamine tetra acetic acid (EDTA) (5 mM, Sigma-Aldrich) to prevent platelet activation, and platelets were obtained after the centrifugation of PRP at 800 × g for 20 min. To extract PEVs, platelet concentrate was activated by thrombin (2 U mL^-1^, Solar bio) in a low-speed shaker at room temperature for 30 min, and was centrifuged at 800 × g for 20 min to obtain a supernatant-enriched PEV solution. Then the supernatant was centrifuged at 100000 × g for 70 min. The vesicle-containing particles were washed in PBS and centrifuged at 100000 × g for 70 min. The particles containing purified vesicles were re-suspended in 200 μL sterilized PBS. Each centrifugation was performed at 4°C. The particle size distribution and zeta potential of PEVs in aqueous solution were then measured using dynamic light scattering (DLS). The morphology of PEVs was observed by transmission electron microscope (TEM). Protein expression of platelet lysates and PEVs was determined by SDS-PAGE and Western blotting.

### Preparation and Characterization of Rapamycin-Loaded PEVs

The insoluble rapamycin was first dissolved in a small amount of dimethyl thioether (DMSO) and incubated with PEVs. The rapamycin-loaded PEVs were separated again by high-speed centrifugation. The size distribution and zeta potential of rapamycin-loaded PEVs were measured by DLS. Furthermore, the properties of drug loading and drug release were determined by high performance liquid chromatography (HPLC). Perform the following liquid phase conditions: chromatographic column C18, mobile phase V (acetonitrile): V (water) = 51:49. The flow rate was 1.0 mL/min, UV detection wavelength was 278 nm, the injection volume was 20 μL, and the column temperature was 27°C. The mass fraction of rapamycin–PEVs was 11.73% when the drug concentration was 100 μg mL^-1^.

### Intranasal Delivery of PEVs

We intranasally delivered rapamycin-PEVs to mice after shallow anesthesia using isoflurane gas containing oxygen, ensuring that the mice recovered within minutes, with 7 mice in each group. We placed each mouse in an air-numbed chamber and adjusted the oxygen flowmeter to 0.6 to 1.2 L/min. Once fully anesthetized, the animals lay on their backs at a 60° angle and a steady rate of respiration was monitored. We then slowly injected 20 μL PEVs into the nostrils at a rate of 5 μL/drop. We stopped the drug for 3 to 4 minutes after each administration to ensure that the mice inhaled the drops and breathed at a steady rate, carefully watching the nostrils for signs of blockage or irritation. After the full dose was given, each mouse recovered from anesthesia before being transferred to its cage.

### Biodistribution of PEVs

Mice were first anesthetized with isoflurane mixed with oxygen, followed by intranasal administration of DiR-labelled PEVs or free DiR, with 4 mice in each group. The IVIS Spectral Imaging System (PerkinElmer Ltd.) was used to monitor NIRF imaging for different groups at different time points over a 24-hour period. Then, major organs were collected for *in vitro* NIRF imaging. Fluorescence intensity was quantified as average radiance (photons sec^-1^ cm^-2^ sr^-1^) with IVIS Living Image 4.2. In addition, brain single-cell suspensions were prepared for flow analysis of PEVs uptake. The expression level of DiR in brain tissue was observed by confocal imaging.

### Histological Analysis

After treatment, major organs of mice in each group were obtained, cleaned in PBS to remove excess blood, fixed in 4% paraformaldehyde solution, and then embedded in paraffin. The paraffin sample was cut into 4 μm thick slices for hematoxylin-eosin staining. Then the pathological status of the samples was observed and analyzed by optical microscopy. In addition, behavioral tests themselves affect the brain, so we performed brain tissue analysis on mice that did not undergo behavioral tests.

### Flow Cytometric Immunoassay of Mouse Tissue

According to the experimental groups, the brain tissue of mice was obtained, and the tissue cell suspension was obtained using a tissue grinder. Then the single cell suspension was removed through the filter screen, washed and centrifuged with PBS, and resuspended in FACS buffer solution (PBS containing 3% BSA). Furthermore, the cells were stained with anti-CD45-PE, anti-CD14-APC (BioLegend). The stained cells were analyzed using BD Accuri C6 flow cytometer and analyzed using Flowjo_V10 software based on 100,000 gated events.

### Statistical Analysis

All data in the present study are means ± standard deviations. The significance of differences between two groups was calculated by a two-tailed unpaired Student’s t test. In addition, analysis of variance (ANOVA) comparisons and Tukey post hoc tests were performed between more than two groups (multiple comparisons). All statistical analyses were performed using GraphPrism (v5.0). Values of *P* = 0.05 or less were considered significant. All intensities of fluorescence expression in the experiments were further calculated by ImageJ software. The standard symbols were presented as \**P* < 0.05, \*\**P*< 0.01, \*\*\**P* < 0.001, and \*\*\*\**P* < 0.0001.

## ASSOCIATED CONTENT

### Supporting Information

The PDF includes Supplementary Figures.

## AUTHOR INFORMATION

### Corresponding Authors

*.

### Author Contributions

C.W. designed the project. Q.M. C. Y., Y. W., H. W. performed the experiments and collected the data. All authors analyzed and interpreted the data, contributed to the writing of the manuscript, discussed the results and implications, and edited the manuscript at all stages.

## ACKNOWLEDGMENTS

This work was supported by the National Key Research and Development Program of China (2022YFB3808100). This work was supported by National Natural Science Foundation of China (No. 32022043). This work is partly supported by Collaborative Innovation Center of Suzhou Nano Science & Technology, the Priority Academic Program Development of Jiangsu Higher Education Institutions (PAPD), the 111 Project.

