## supplementary figures for "Neurological Disorder after Severe Pneumonia is Associated with Translocation of Bacteria from Lung to Brain"

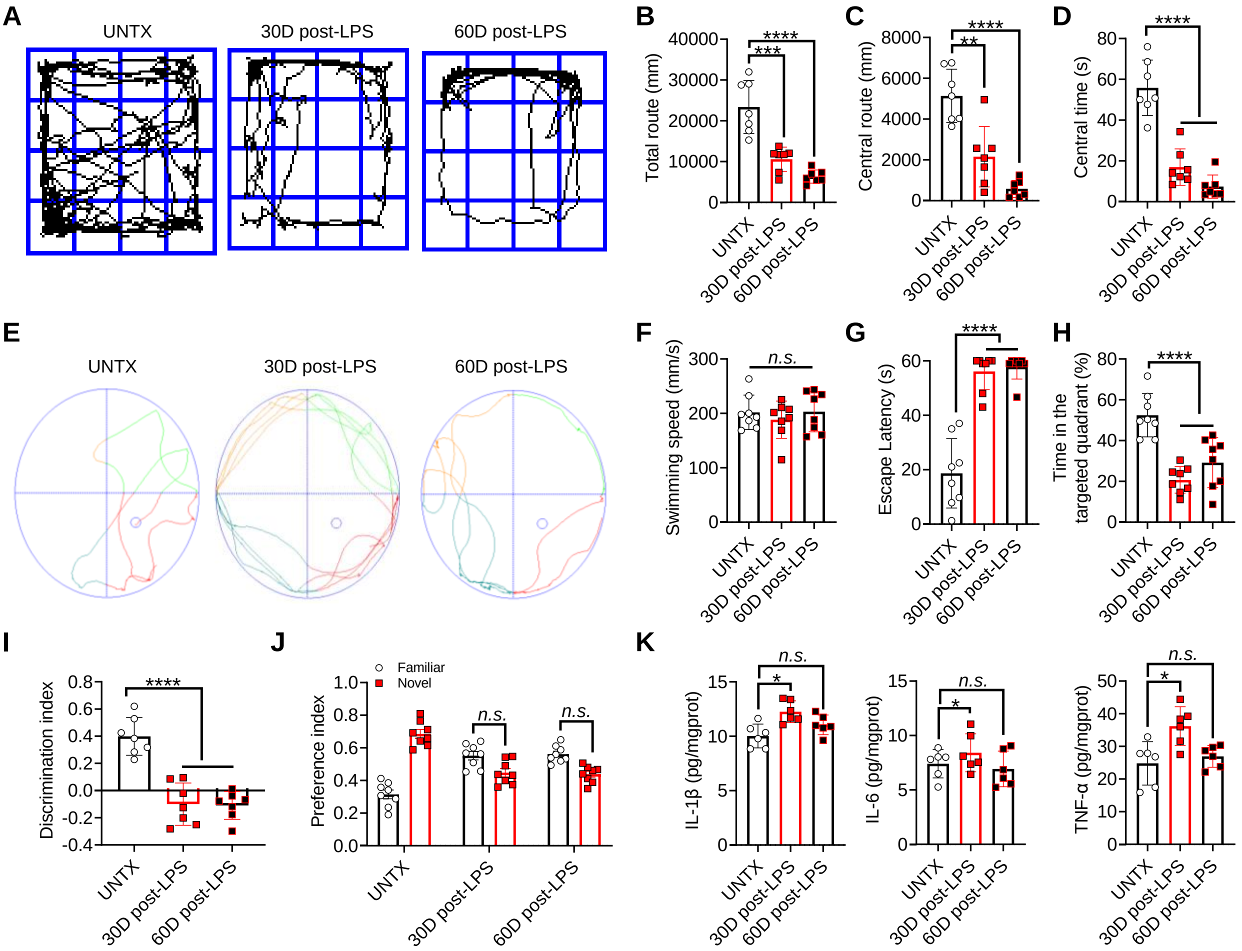


**Supplementary Figure S1.** (**A**) The representative paths of mice in the open field test and quantitative analysis of (**B**) the total route, (**C**) the central route and (**D**) the central time. (**E**) The representative plot of Morris water maze images and quantification of (**F**) swimming speed, (**G**) escape latency, and (**H**) the ratio of the time in the target quadrant to the total time. (**I**) Discrimination index in the novel object recognition. (**J**) Preference index in the novel object recognition. (**K**) Inflammatory factors including IL-1β, IL-6, and TNF-α of brain tissue homogenate. Data are shown as mean ± SD (n=6-8). Statistical significance was calculated by Student's t-test (two-tailed) and one-way ANOVA using the Tukey posttest. *$P < 0.05$; **$P< 0.01$; ***$P < 0.001$; ****$P < 0.001$; n.s.; nonsignificant.

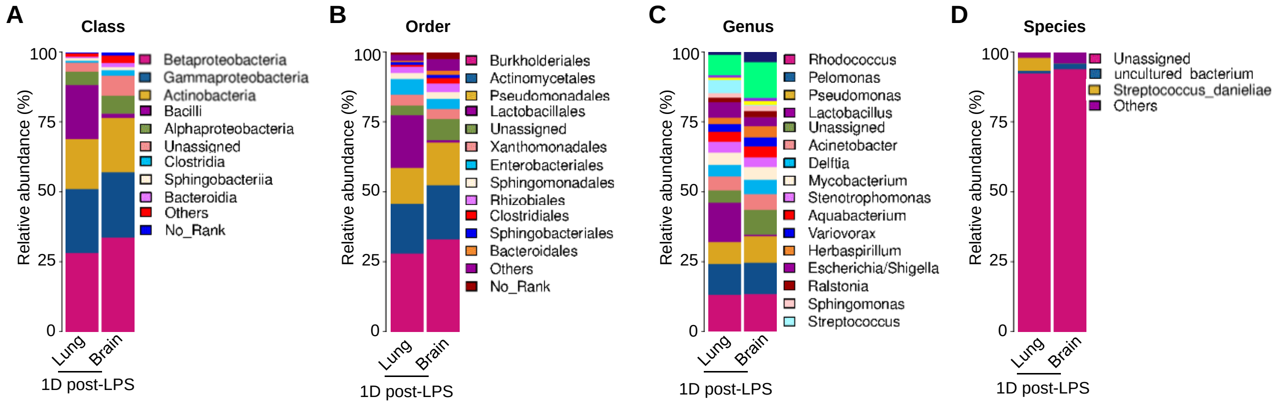


**Supplementary Figure S2.** (**A-D**) Relative abundance of lung and brain bacterial inhabitants at the class (**A**), order (**B**), genus (**C**), species (**D**) level.

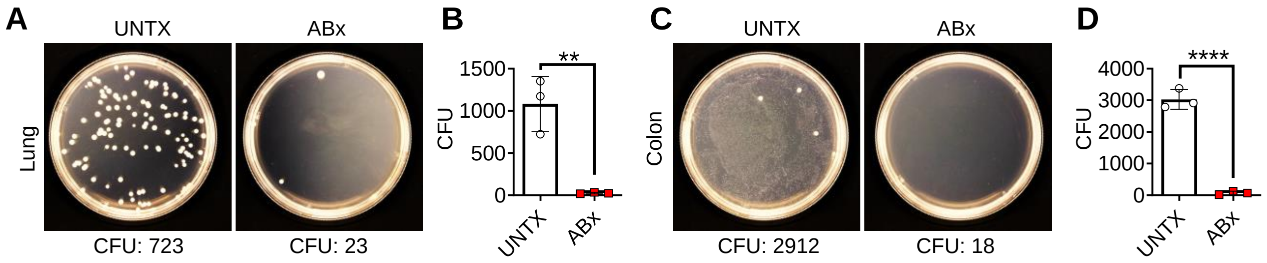


**Supplementary Figure S3.** (**A**) The representative plot of bacterial colony growth after 24 hours of lung tissue homogenate and (**B**) corresponding quantification results of colony forming units. (**C**) The representative plot of bacterial colony growth after 24 hours of colon tissue homogenate and (**D**) corresponding quantification results of colony forming units. Data are shown as mean ± SD (n=3). Statistical significance was calculated by Student's t-test (two-tailed) and one-way ANOVA using the Tukey posttest. **$P$< 0.01; ****$P$ < 0.001.

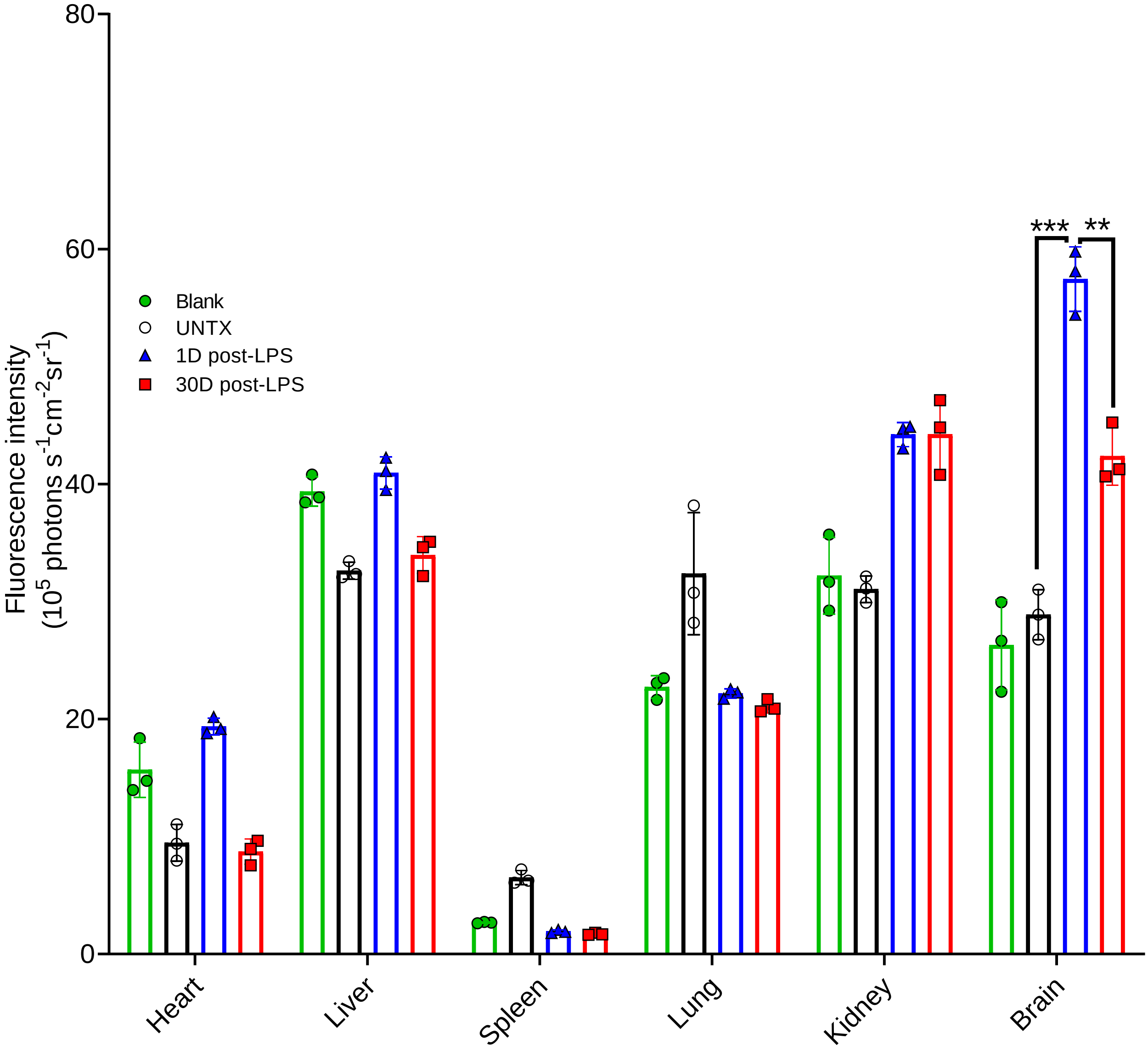


**Supplementary Figure S4.** Quantification result of biodistribution of 4-kDa FITC–conjugated dextran. Data are shown as mean ± SEM (n=3). Statistical significance was calculated by Student's t-test and one-way ANOVA using the Tukey posttest. ***P* < 0.01; ****P* < 0.001.

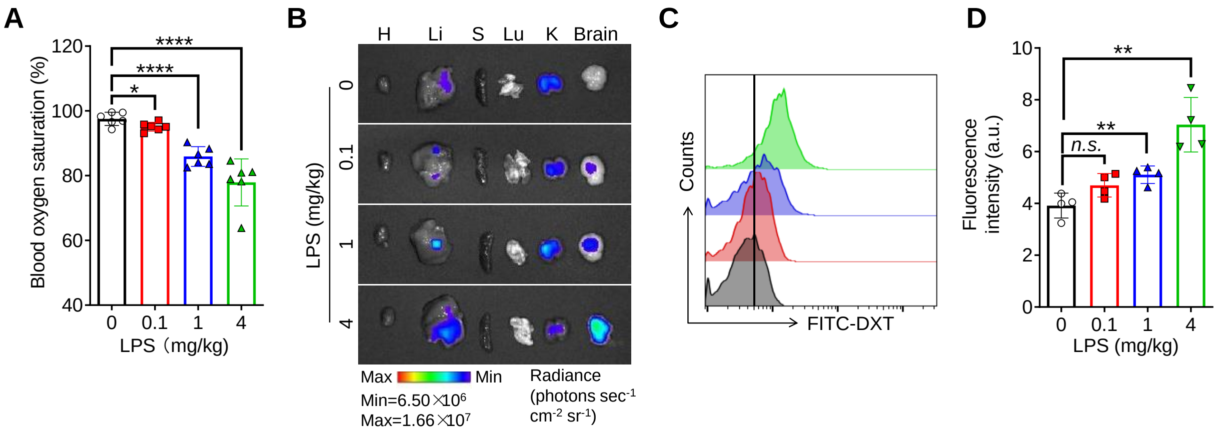


**Supplementary Figure S5.** (**A**) Blood oxygen saturation data of mice after different treatments as indicated. (**B**) *Ex vivo* imaging showed biodistribution of 4-kDa FITC–conjugated dextran. (**C**) Representative flow cytometric analysis of 4-kDa FITC–conjugated dextran signal and (**D**) corresponding quantification results of MFI of FITC. Data are shown as mean ± SD (n=4-6). Statistical significance was calculated by Student’s t-test (two-tailed) and one-way ANOVA using the Tukey posttest. *$P < 0.05$; **$P< 0.01$; ****$P < 0.001$; n.s.; nonsignificant.

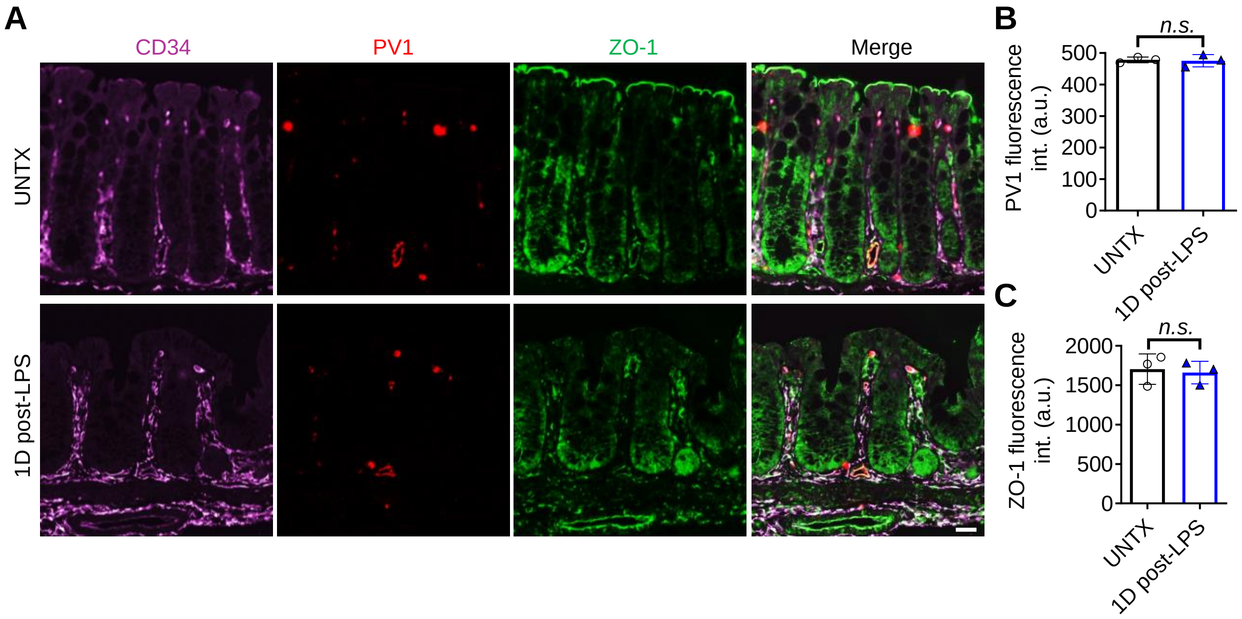


**Supplementary Figure S6.** (**A**) Representative confocal images showing PV1 (red) detection in CD34$^+$ (purple) blood vessels and ZO-1 (green) in colon of mice after different treatments as indicated, scale bars, 10 µm, and (**B**, **C**) corresponding quantitative analysis. Data are shown as mean ± SD (n=3). Statistical significance was calculated by Student's t-test (two-tailed) and one-way ANOVA using the Tukey posttest. n.s.; nonsignificant. a. u., arbitrary units.

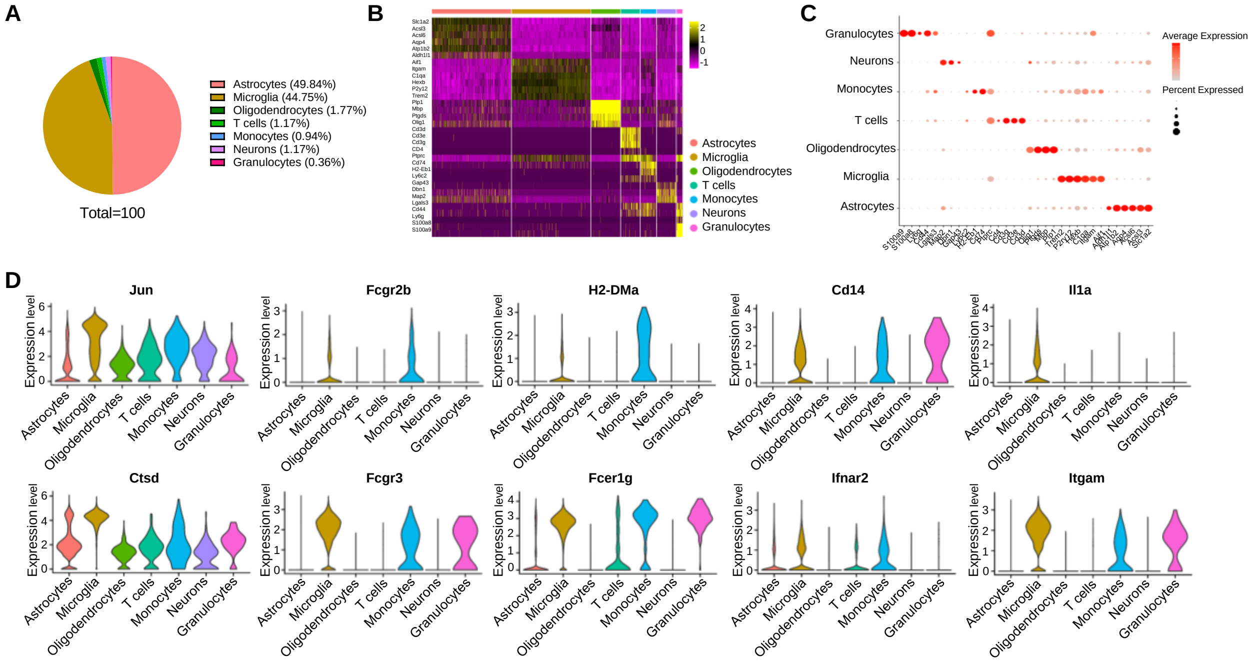


**Supplementary Figure S7.** (**A**) The percentage of each cluster. (**B**) Heatmap displaying of marker genes expression in all brain cell clusters. (**C**) Dotplot of marker genes expression in all brain cell clusters. (**D**) Violin diagram of gene expression in brain cells.

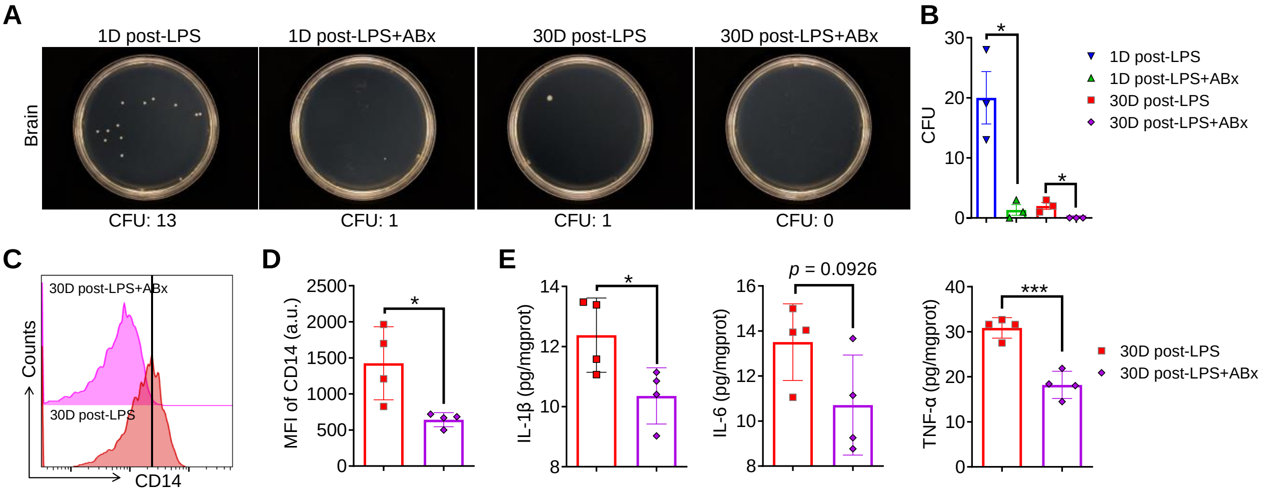


**Supplementary Figure S8.** (**A**) The representative plot of bacterial colony growth after 24 hours of brain tissue homogenate and (**B**) corresponding quantification results of colony forming units. (**C**) Representative flow cytometric analysis of CD14 and (**D**) corresponding quantification results of MFI of CD14. (**E**) Inflammatory factors including IL-1β, IL-6, and TNF-α of brain tissue homogenate. Data are shown as mean ± SD (n=3-5). Statistical significance was calculated by Student's t-test (two-tailed) and one-way ANOVA using the Tukey posttest. *$P < 0.05$; ***$P < 0.001$; n.s.; nonsignificant.

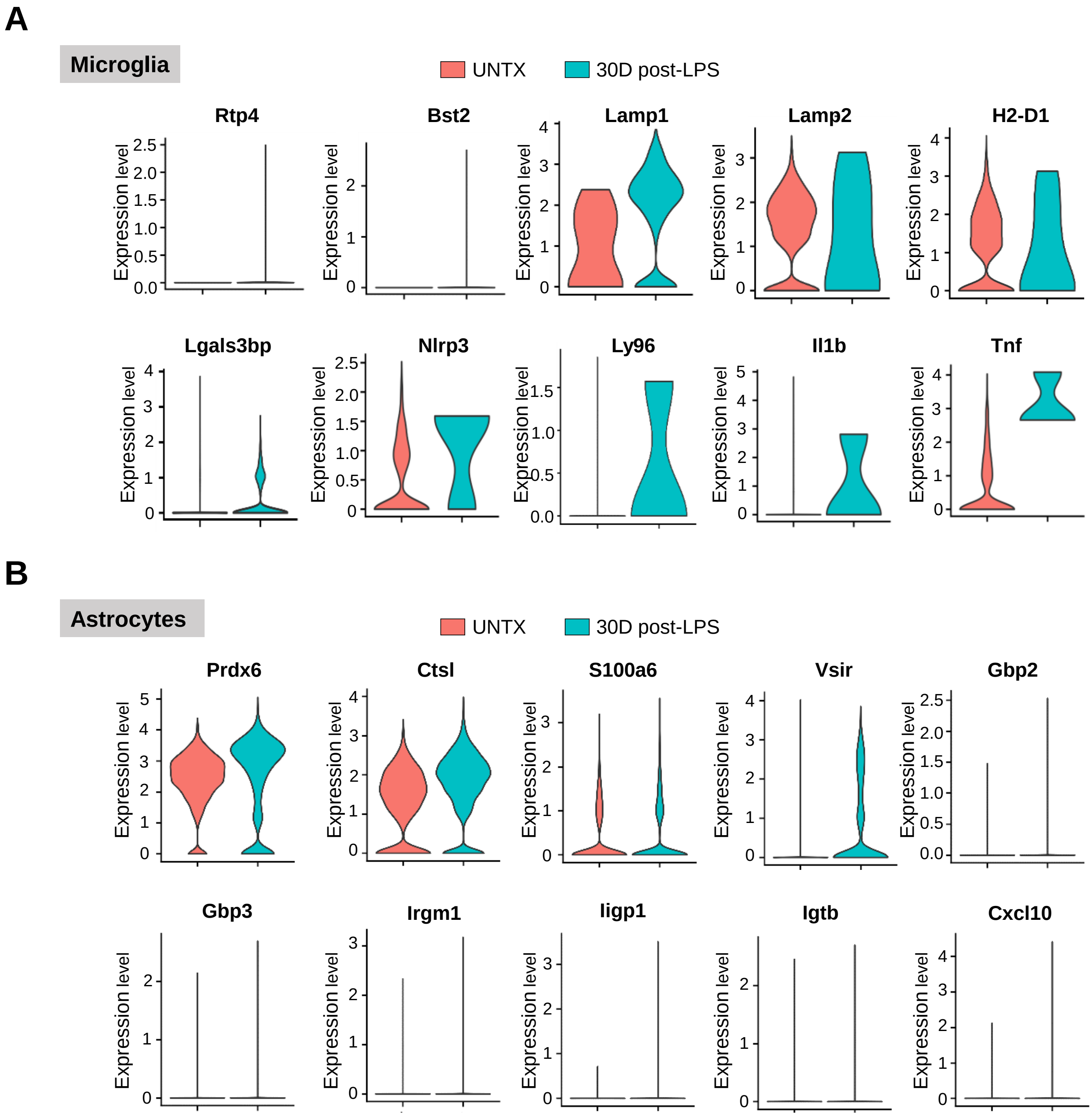


**Supplementary Figure S9.** (**A**) Violin diagram of gene expression in microglia. (**B**) Violin diagram of gene expression in astrocytes.

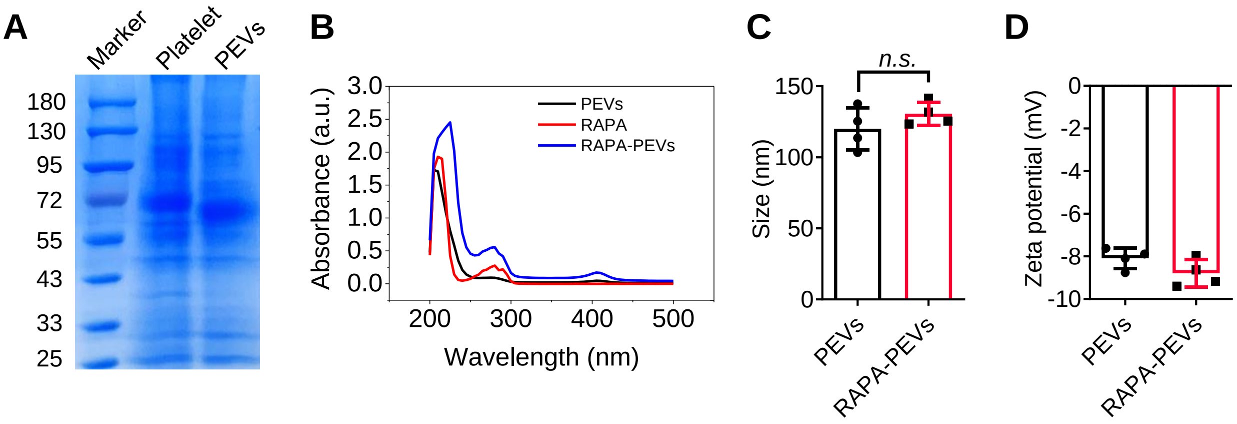


**Supplementary Figure S10.** (**A**) SDS-PAGE of platelet lysate and PEVs with Coomassie brilliant blue staining. (**B**) Representative UV–vis absorption peaks of PEVs, RAPA, RAPA-PEVs in the phosphate-buffered saline. (**C**) The size and (**D**) the zeta potential of the PEVs and RAPA-loaded PEVs. Data are shown as mean ± SD (n=4). Statistical significance was calculated by Student's t-test (two-tailed) and one-way ANOVA using the Tukey posttest. n.s.; nonsignificant.

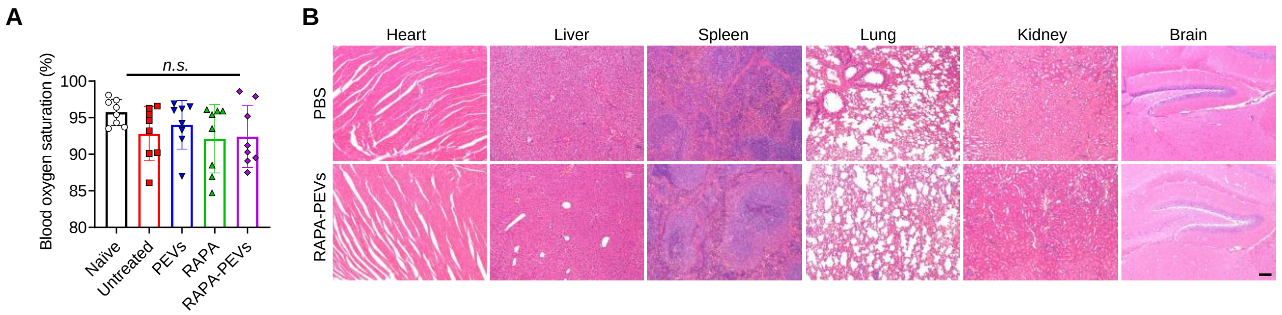


**Supplementary Figure S11.** (**A**) Blood oxygen saturation data of mice after different treatments as indicated. (**B**) Representative data for hematoxylin and eosin staining in major organs from WT mice treated with PBS, or RAPA-PEVs. Scale bars, 100 μm. Data are shown as mean ± SD (n=8). Statistical significance was calculated by Student's t-test (two-tailed) and one-way ANOVA using the Tukey posttest. n.s.; nonsignificant.

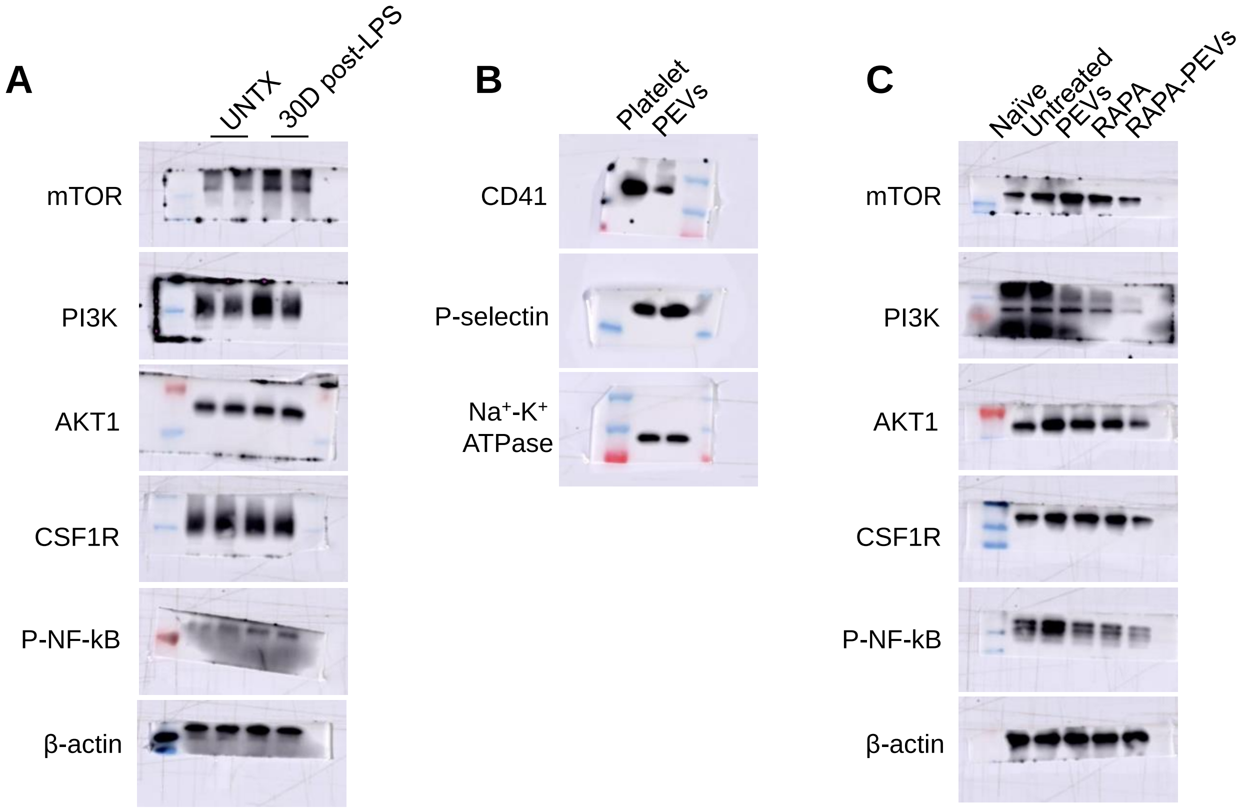


**Supplementary Figure S12.** Western blot analysis of the expression of various types of proteins after various treatments as indicated.

| Reagent or Resource | Source | Identifier |
|---|---|---|
| **Antibodies** | | |
| APC anti-mouse CD14 | Biolegend | Cat# 123312 |
| PE anti-mouse CD45 | Biolegend | Cat# 103106 |
| Anti-mTOR | Servicebio | Cat# GB111839 |
| Anti-PI3k | Abclonal | Cat# A4992 |
| Anti-AKT1 | Servicebio | Cat# GB111114 |
| Anti-CSF1R | Abclonal | Cat# A3019 |
| Anti-P-NF-kB | Abclonal | Cat# AP0475 |
| Anti-β-actin | Servicebio | Cat# GB11001 |
| Anti-Aβ | Servicebio | Cat# GB111197 |
| Anti-IBA-1 | Servicebio | Cat# GB11105 |
| Anti-GFAP | Servicebio | Cat# GB11096 |
| Anti-PV-1 | Servicebio | Cat# GB113141 |
| Anti-ZO-1 | Servicebio | Cat# GB111402 |
| Anti-CD31 | Servicebio | Cat# GB110632 |
| Anti-CD34 | Servicebio | Cat# GB111693 |
| Goat anti-rabbit IgG-HRP | Absin | Cat# abs20040 |
| **Drug** | | |
| Rapamycin | Energy chemical | Cat# E080201 |
| LPS | Biosharp | Cat# bs904 |
| Ampicillin | Aladdin | Cat# A105484 |
| Gentamicin | Aladdin | Cat# G100391 |
| Metronidazole | Energy Chemical | Cat# A040096 |
| Vancomycin | Aladdin | Cat# V105495 |
| **Critical Commercial Assays** | | |
| BCA Protein Assay Kit | Beyotime | Cat# P0012 |
| PAGE Gel Rapid Preparation Kit | Epizyme | Cat# PG113 |
| Commassie Blue Staining Solution | Epizyme | Cat# PS111 |
| Mouse IL-1 beta Uncoated ELISA Kit | Invitrogen | Cat# 88-7013-88 |
| Mouse IL-6 Uncoated ELISA Kit | Invitrogen | Cat# 88-7064-88 |
| Mouse TNF alpha Uncoated ELISA Kit | Invitrogen | Cat# 88-7324-88 |

**Table. S1 Materials used in study.**
